# SIEVE: joint inference of single-nucleotide variants and cell phylogeny from single-cell DNA sequencing data

**DOI:** 10.1101/2022.03.24.485657

**Authors:** Senbai Kang, Nico Borgsmüller, Monica Valecha, Jack Kuipers, Joao Alves, Sonia Prado-López, Débora Chantada, Niko Beerenwinkel, David Posada, Ewa Szczurek

## Abstract

Single-cell DNA sequencing (scDNA-seq) has enabled the identification of single nucleotide somatic variants and the reconstruction of cell phylogenies. However, statistical phylogenetic models for cell phylogeny reconstruction from raw sequencing data are still in their infancy. Here we present SIEVE (SIngle-cell EVolution Explorer), a statistical method for the joint inference of somatic variants and cell phylogeny under the finite-sites assumption from scDNA-seq reads. SIEVE leverages raw read counts for all nucleotides at candidate variant sites, and corrects the acquisition bias of branch lengths. In our simulations, SIEVE outperforms other methods both in phylogenetic accuracy and variant calling accuracy. We apply SIEVE to three scDNA-seq datasets, for colorectal (CRC) and triple-negative breast cancer (TNBC), one of them generated by us. On simulated data, SIEVE reliably infers homo-and heterozygous somatic variants. The analysis of real data uncovers that double mutant genotypes are rare in CRC but unexpectedly frequent in TNBC samples.

## Introduction

Intra-tumour heterogeneity is a consequence of accumulated somatic mutations during tumour evolution [1, 2] and the culprit of acquired resistance and relapse in clinical cancer therapy [3, 4]. Phylogenetic inference is a powerful tool to understand the development of intra-tumour heterogeneity in time and space. Variant allele profiles derived from bulk sequencing data have typically been used to reconstruct the tumour phylogeny at the level of clones [5–9]. More recently, the development of scDNA-seq [10–12] has enabled single-nucleotide variant (SNV) calling [13–18] and phylogeny reconstruction [15, 19–26] down to the single-cell level.

A statistical phylogenetic model is defined by an instantaneous transition rate matrix, a tree topology and tree branch lengths. Such a model defines a Markov process for the evolution of nucleotides or genotypes [27]. Studying the evolutionary process and estimating important parameters such as the branch lengths using statistical phylogenetic models has a long tradition, benefits from well established theory, and has many applications, such as interpreting temporal cell dynamics [28].

However, compared to statistical phylogenetic models, most methods for phylogeny reconstruction from scDNA-seq operate within a simpler modelling framework. First, although branch lengths are a critical part of a phylogenetic tree and reflect the real evolutionary distances among cells, they are often ignored. Those approaches that do infer branch lengths [22, 26] employ the data from the variant sites and ignore information from *background sites* (that have a wildtype genotype), which may lead to so-called acquisition bias and overestimated branch lengths [29, 30].

Moreover, variant calling and phylogenetic inference are commonly considered independent tasks. Variant calling is typically performed first, and phylogenetic inference is performed on the called variants. However, variant calling, particularly from scDNA-seq data, can be hampered by missing data and low coverage, potentially resulting in wrong calls that could mislead phylogenetic inference. A feasible strategy to alleviate this problem is to integrate tree reconstruction with variant calling [12], where phylogenetic information on cell ancestry is used to obtain more reliable variant calls. Recently developed methods for scDNA-seq data approach this strategy from different perspectives [15, 31]. However, those methods do not operate within the statistical phylogenetic framework, in particular do not infer branch lengths of the tree. Moreover, either they fully follow the infinite-sites assumption (ISA), which is often violated in real datasets [32, 33], or relax this assumption to only a limited extent. As a result, they may miss important events in the evolution of tumours. Thus, methods have not yet been developed which, employing statistical phylogenetic models under the finite-sites assumption (FSA), infer cell phylogeny from raw scDNA-seq data and simultaneously call variants.

To address this, we propose SIEVE, a statistical method that exploits raw read counts for all nucleotides from scDNA-seq to reconstruct the cell phylogeny and call variants based on the inferred phylogenetic relations among cells. To our knowledge, SIEVE is the first approach that employs a statistical phylogenetic model following FSA, where branch lengths, measured by the expected number of somatic mutations per site, are corrected for the acquisition bias using the data from the background sites, and simultaneously calls variants and allelic dropout (ADO) states from raw read counts data. The model is able to detect twelve different types of mutation events in evolutionary history. SIEVE is implemented and available as a package of BEAST 2, which allows for benefitting from other packages in this framework. Using simulated data, we assess the performance of our model in comparison to existing methods. To illustrate the functionality of SIEVE, we apply it to datasets from two patients with CRC and one with TNBC.

## Results

### SIEVE is a statistical method for joint inference of SNVs and cell phylogeny from scDNA-seq data

SIEVE takes as input raw read count data at candidate SNV sites, accounting for the read counts for three alternative nucleotides and the total depth at each site (Fig. 1a) and combines a statistical phylogenetic model with a probabilistic graphical model of the read counts, incorporating a Dirichlet Multinomial distribution of the nucleotide counts (Fig. 1b; Methods). The statistical phylogenetic model allows for acquisition and loss of mutations on both maternal and paternal alleles (Fig. 1c). It considers four possible genotypes, 0/0 (referred to as *wildtype*), 0/1 (*single mutant*), 1/1 (*double mutant*, where the two alternative nucleotides are the same) and 1/1^′^ (*double mutant*, where the two alternative nucleotides are different). With these genotypes, SIEVE is able to discern twelve different types of mutation events (Table 2; Methods). Based on the inferred tree (Fig. 1d), SIEVE calls the maximum likelihood somatic mutations (Fig. 1e). The tree contains a trunk joining the root representing a healthy cell with the most recent common ancestor (MRCA) of the modelled cells, representing the acquisition of clonal mutations at the initial stage of tumour progression. SIEVE leverages the noisy raw read counts to integrate genotype uncertainty into cell phylogeny inference. Benefiting from the inferred cell relationships, SIEVE is able to reliably infer the single-cell genotypes, especially for sites where only few reads are available. SIEVE is implemented as a package of BEAST 2, a flexible and mature framework for statistical phylogenetic modelling [34].

**Fig. 1:**
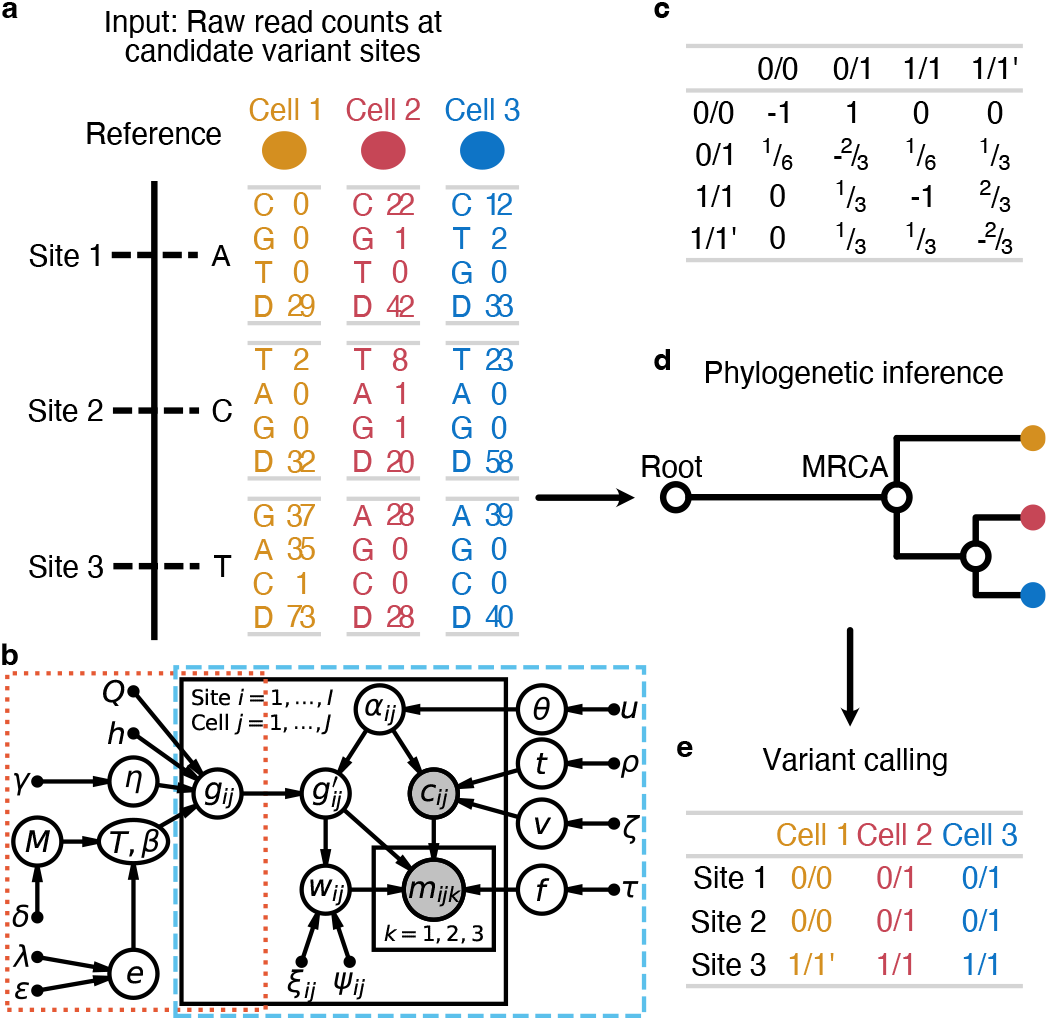
Overview of the SIEVE model. **a**, Input data to SIEVE at candidate SNV sites. For a specific cell at an SNV site, fed to SIEVE are the read counts for all nucleotides: reads of the three alternative nucleotides with values in descending order and the total coverage (denoted by D in **a**). **b**, Graphical representation of the SIEVE model. Bridged by *g*_*ij*_, the genotype for site *i* in cell *j*, the orange dotted frame encloses the statistical phylogenetic model, and the blue dashed frame highlights the model of raw read counts. Shaded circle nodes represent observed variables, while unshaded circle nodes represent hidden random variables. Small filled circles correspond to fixed hyper parameters. Arrows denote local conditional probability distributions of child nodes given parent nodes. The sequencing coverage *c*_*ij*_ follows a negative binomial distribution parameterised by the number of sequenced alleles *α*_*ij*_, the mean of allelic coverage *t* and the variance of allelic coverage *v. α*_*ij*_ is a hidden categorical variable parameterised by ADO rate *θ*, which has a uniform prior with fixed hyper parameter *u. t* also has a uniform prior with fixed parameter *ρ*, while *v* has an exponential prior parameterised by *ζ*. The nucleotide read counts ***m***_***ij***_ given *c*_*ij*_ follow a Dirichlet-multinomial distribution parameterised by ADO-affected genotype 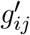, which is a hidden random variable depending on *α*_*ij*_ and genotype *g*_*ij*_, effective sequencing error rate *f*, which has en exponential prior with fixed hyper parameter *τ*, and overdispersion *w*_*ij*_, which is a hidden categorical variable dependent on 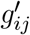 parameterised by fixed parameters *ξ*_*ij*_ and *ψ*_*ij*_ for each category. *g*_*ij*_ is determined by the statistical phylogenetic model parameterised by fixed rate matrix *Q*, fixed number of categories *h* as well as shape parameter *η* with exponential prior for site-wise substitution rates, and tree topology *𝒯* along with branch lengths ***β***. *𝒯* and ***β*** have a coalescent prior with an exponentially growing population parameterised by effective population size *M*, which has a multiplicative inverse prior, and growth rate *e*, which has a laplace prior parameterised by *λ* and *ϵ*. **c**, The transition rate matrix in the statistical phylogenetic model. During an infinitesimal time interval only one change is allowed to occur. **d**, The cell phylogeny inferred from the data with SIEVE. Not only is the tree topology crucial, but also the branch lengths. The root represents a normal cell, and the only direct child of the root is the most recent common ancestor (MRCA) of all cells. **e**, Variant calling given the inferred cell phylogeny. For further details see Methods.

We investigated the performance of SIEVE using simulated data with different means and variances of allelic coverage, reflecting different *coverage qualities* (Methods). Specifically, we simulated data with low mean and high variance of allelic coverage (low quality), with high mean and medium variance (medium quality), and with high mean and low variance (high quality). Other important dataset characteristics were varied, including the number of cells and mutation rate, which is measured by the number of accumulated somatic mutations per site per generation.

### SIEVE accurately estimates tree topology and branch lengths

We first evaluated the accuracy of SIEVE in inferring the simulated cell phylogeny with branch lengths using the rooted branch score (BS) distance [35] (Fig. 2a and Methods). We compared to CellPhy [26] and SiFit [22], which were fed with the variant calls from Monovar [13]. Here, we gave SiFit an advantage of setting the true positive error rate used in the simulation (Methods). Thanks to the acquisition bias correction, SIEVE reports branch lengths as expected number of somatic mutations per site, while CellPhy and SiFit per SNV site. SCIPhI [15] does not infer branch lengths, hence its rooted BS distance could not be computed. SIEVE consistently outperformed CellPhy and SiFit, regardless of the number of cells, mutation rate and coverage quality. This may be because, in contrast to SIEVE, CellPhy and SiFit do not model raw reads and, importantly for the rooted BS distance, do not correct the inferred branch lengths for acquisition bias. We also found that the rooted BS distance of SIEVE had a negative nonlinear association with the number of background sites (Extended Data Fig. 1), explaining the relatively greater differences under higher mutation rates. These results proved the necessity for correcting the acquisition bias with enough background sites to obtain accurate branch lengths.

**Fig. 2:**
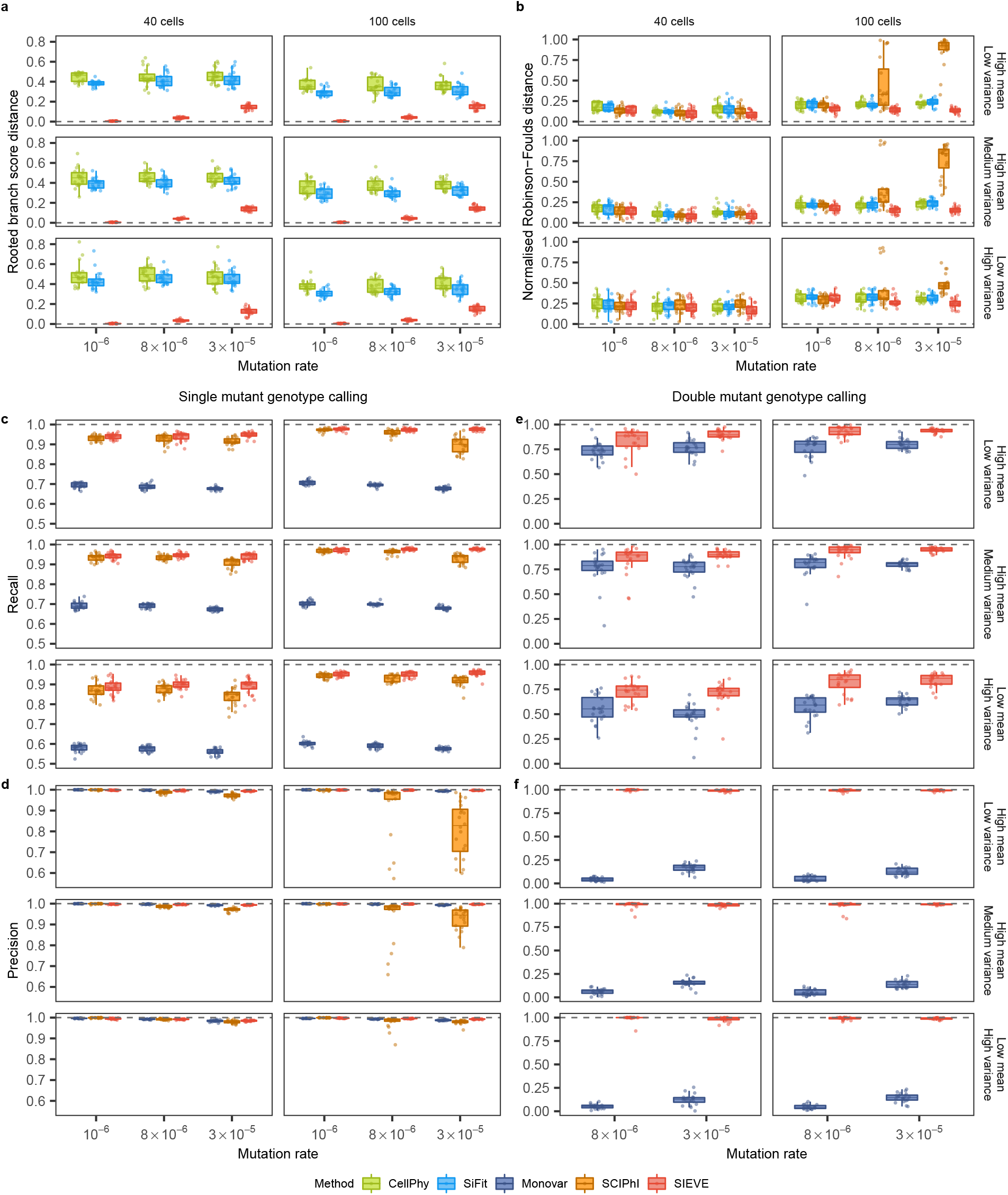
Benchmarking result of the SIEVE model. Varying are the number of tumour cells, mutation rate and coverage quality. Each simulation is repeated *n* = 20 times with each repetition denoted by coloured dots. The grey dashed lines represent the optimal values of each metric. Box plots comprise medians, boxes covering the interquartile range (IQR), and whiskers extending to 1.5 times the IQR below and above the box. **a-b**, Box plots of the tree inference accuracy measured by the rooted BS distance where the branch lengths are taken into account (**a**) and the normalised RF distance where only tree topology is considered (**b**). **c-d**, Box plots of the single mutant genotype calling results measured by the fraction of true positives respectively in the ground truth positives, i.e., the sum of true positives and false negatives, (recall, **c**) as well as in the predicted positives, i.e., the sum of true positives and false positives, (precision, **d**). **e-f**, Box plots of the double mutant genotype calling results measured by recall (**e**) and precision (**f**), where the variant calling results when mutation rate is 10^−6^ are omitted as very few double mutant genotypes are generated (less than 0.1%).

As the rooted BS distance is dominated by the branch lengths, we further assessed SIEVE’s accuracy in inferring the tree structure using the normalised Robinson-Foulds (RF) distance [36]. Compared to CellPhy, SiFit and SCIPhI (Fig. 2b and Methods), SIEVE was the most robust method to changes of mutation rate, number of cells and coverage quality. When the data hardly contained mutations violating the ISA (mutation rate being 10^−6^, with less than 0.1% double mutant genotypes and at most 1% SNV sites with parallel mutations), all methods achieved a similar median RF distance (around 0.15-0.3). Since in contrast to SCIPhI, SIEVE, CellPhy and SiFit employ statistical phylogenetic models following FSA, this indicates that models following FSA are also applicable to data evolving under the ISA. SIEVE outperformed CellPhy and SiFit when the number of cells and the mutation rate increased. When the data clearly violated the ISA (mutation rates being 8 × 10^−6^ and 3 × 10^−5^, with 0.02%-0.3% and 0.1%-1% double mutant genotypes, as well as 2%-8% and 10%-27% SNV sites with parallel mutations indicative of FSA, respectively), SCIPhI inferred reasonable tree topologies from datasets with a small number of cells (40). However, its performance dramatically dropped with 100 cells, especially when the data was of medium or high coverage quality. The behaviour of SCIPhI might be related to its estimation of ADO rate and single mutant genotype calling in these scenarios.

### SIEVE accurately infers parameters in the model of raw read counts

We next investigated the accuracy of parameter estimates, including *effective* sequencing error rate, ADO rate, and wildtype and alternative overdispersion (Extended Data Fig. 2 and Methods). Here, the effective sequencing error rate (Extended Data Fig. 2a) takes into account both amplification and sequencing error rates in scDNA-seq. Wildtype and alternative overdispersion are parameters in the distribution of nucleotide read counts related to different genotypes. The former corresponds to genotype 0/0 and 1/1, while the latter to genotype 0/1 and 1/1^′^. SIEVE accurately inferred most parameters in all simulated scenarios regardless of the number of cells, mutation rate and coverage quality. Although SIEVE’s accuracy of estimating ADO rate slightly decreased with the coverage quality, it still was the best among the competing methods. For data with medium and high coverage quality, 100 cells and higher mutation rates (8 × 10^−6^ and 3 × 10^−5^), SCIPhI tended to overestimate ADO rates.

### SIEVE accurately calls single and double mutations

Next, we assessed SIEVE’s performance in calling the single mutant genotype (Fig. 2c,d, Extended Data Fig. 3a,b, Extended Data Fig. 4, and Methods). As opposed to Monovar, recall for SIEVE and SCIPhI increased with the number of cells but was less sensitive to the coverage quality (Fig. 2c). The recall of SIEVE was higher than that of SCIPhI by 0.16%-18.55% and that of Monovar by 28.89%-71.74%. Unlike Monovar, both SIEVE and SCIPhI benefit from the information provided by cell phylogenies. We speculate that the advantage of SIEVE over SCIPhI stems from the use of raw read counts for all nucleotides, while SCIPhI only employs the sequencing coverage and the read count of the most prevalent alternative nucleotide.

Moreover, SIEVE and Monovar achieved comparable precision (Fig. 2d) and false positive rates (Extended Data Fig. 3a) regardless of the number of cells, mutation rate and coverage quality. However, this did not hold for SCIPhI. By analysing the types of false positives among the predicted single mutant genotypes (Extended Data Fig. 4 and Methods), we found that SCIPhI tended to miscall wildtype genotypes as single mutant genotype (i.e., 0/0 are called as 0/1) (Extended Data Fig. 4a). This occurred with high mutation rates (8 × 10^−6^ and 3 × 10^−5^), especially in scenarios where SCIPhI inferred inaccurate trees (Fig. 2b) and overestimated ADO rates (Extended Data Fig. 2b). The reason is twofold. First, the ISA upon which SCIPhI builds naturally limits its application to data following FSA. Second, under these scenarios, SCIPhI tends to mistake sites with no variant support for ADO events, and hence its high ADO rate. SIEVE avoids such mistakes by leveraging a model of sequencing coverage (Methods), thereby accounting for the related overdispersion and correctly estimating the ADO rate. We also noticed that when data clearly violated ISA, both Monovar and SCIPhI miscalled more double mutant genotypes as the single mutant genotype than SIEVE (Extended Data Fig. 4b).

We then focused on the results of double mutant genotype calling (Fig. 2e,f, Extended Data Fig. 3c,d and Methods), where SCIPhI was excluded as it is unable to call such mutations. The recall of double mutant genotypes for SIEVE and Monovar increased with the number of cells and the coverage quality (Fig. 2e), while SIEVE showed higher recall for such genotypes than Monovar. Moreover, SIEVE outperformed Monovar with high precision (almost 1, Fig. 2f) and low false positive rate (almost 0, Extended Data Fig. 3c).

### SIEVE accurately calls ADOs for data of adequate coverage quality

We further assessed SIEVE’s performance in ADO calling (Extended Data Fig. 5), where there are no published methods for us to compare with. When calling ADOs, SIEVE’s performance was independent of the number of cells or mutation rate, but highly dependent on the coverage quality. The reason is that SIEVE calls ADOs by inferring the number of sequenced alleles, assuming it is proportional to the observed sequencing coverage (see Methods). Consequently, for data with medium and high coverage quality the average F1 score of ADO calling was high (0.86 and 0.93, respectively), whereas for data with low coverage quality, which is typical for current scDNA-seq data, the ADO calling performance deteriorated, with average F1 score being only 0.10. Since the coverage quality of real data is low, we do not report ADO calling results for all real datasets analysed below (Extended Data Table 1).

### SIEVE inferred a phylogenetic tree and called variants for CRC cells

We applied SIEVE to a new single-cell whole genome sequencing (scWGS) dataset, where 28 tumour cells were isolated from three primary tumour biopsies of a patient with CRC (CRC28; see Methods). We identified 8,470 candidate SNV sites and 1,163,335,103 background sites. To take into account branch-wise substitution rate variation, we employed a relaxed molecular clock model [37] (same for the following datasets; see Methods). In the inferred maximum clade credibility (MCC) tree (Fig. 3; see Extended Data Fig. 6 for the branch lengths), tumour cells grouped into three highly supported clades corresponding to the three biopsies. The estimated effective sequencing error and ADO rates were 7.6 × 10^−4^ and 0.20, respectively.

**Fig. 3:**
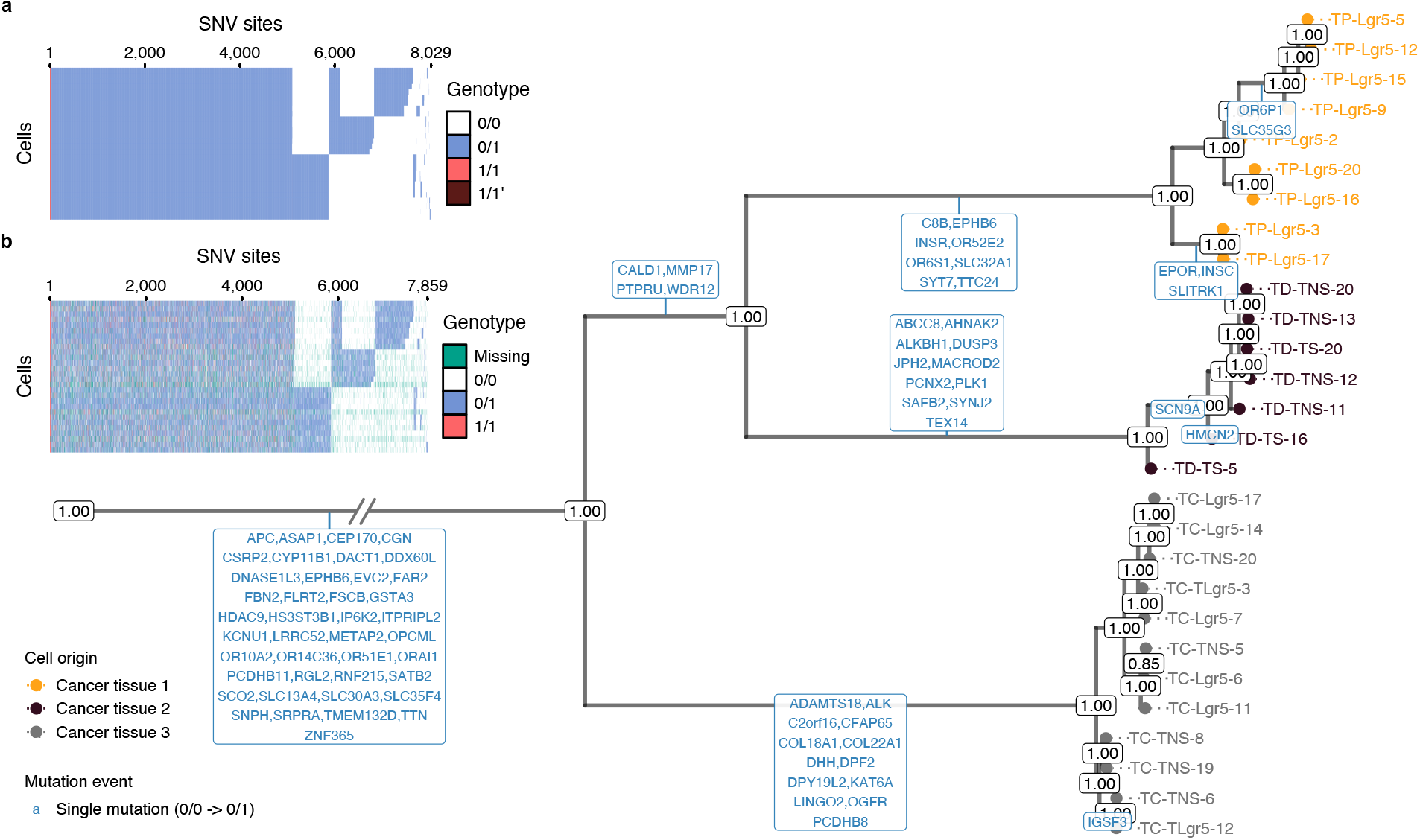
Results of phylogenetic inference and variant calling for CRC28 dataset. Shown is SIEVE’s maximum clade credibility tree. The exceptionally long trunk has been folded (marked by slashes). Cells are coloured according to the corresponding biopsies. The numbers at each node represent posterior probabilities (threshold *p >* 0.5). At each branch, genes with non-synonymous mutations are depicted in blue. **a-b**, Variant calling heatmap for SIEVE (**a**) and Monovar (**b**). Listed in the legend are the categories of predicted genotypes by each method. Cells in the row are in the same order as that of leaves in the phylogenetic tree.

We mapped non-synonymous mutations to the internal branches (Methods), where only single mutations were found, indicating that the evolution of these mutational process likely followed the ISA. Many mutations resided on the trunk (clonal mutations), including established CRC driver genes [38, 39], such as *APC*.

SIEVE identified 8,029 SNV sites among the candidate SNV sites (Fig. 3a), where most of the genotypes were single mutant and few were double mutant, including 1/1^′^. The variant calling results of SIEVE and Monovar (Fig. 3b) were overall similar. However, the calls from Monovar were clearly more noisy, with many missing entries and more double mutant genotypes, some of which might be false positives according to the simulation results. The proportion of genotypes called by SIEVE and Monovar were summarised in Supplementary Table 1 (same for the following datasets).

### SIEVE inferred a phylogenetic tree and called variants for TNBC cells

We then applied SIEVE to a single-cell whole exome sequencing (scWES) dataset [40], containing 16 tumour cells collected from a patient with TNBC (TNBC16; see Methods). We identified 5,912 candidate SNV sites and 152,027,822 background sites. The estimated tree was supported by high posterior probabilities (Fig. 4) with a relatively long trunk and short terminal branches (Extended Data Fig. 7). We estimated that the effective sequencing error rate was 8.2 × 10^−4^ and the ADO rate was 0.05.

**Fig. 4:**
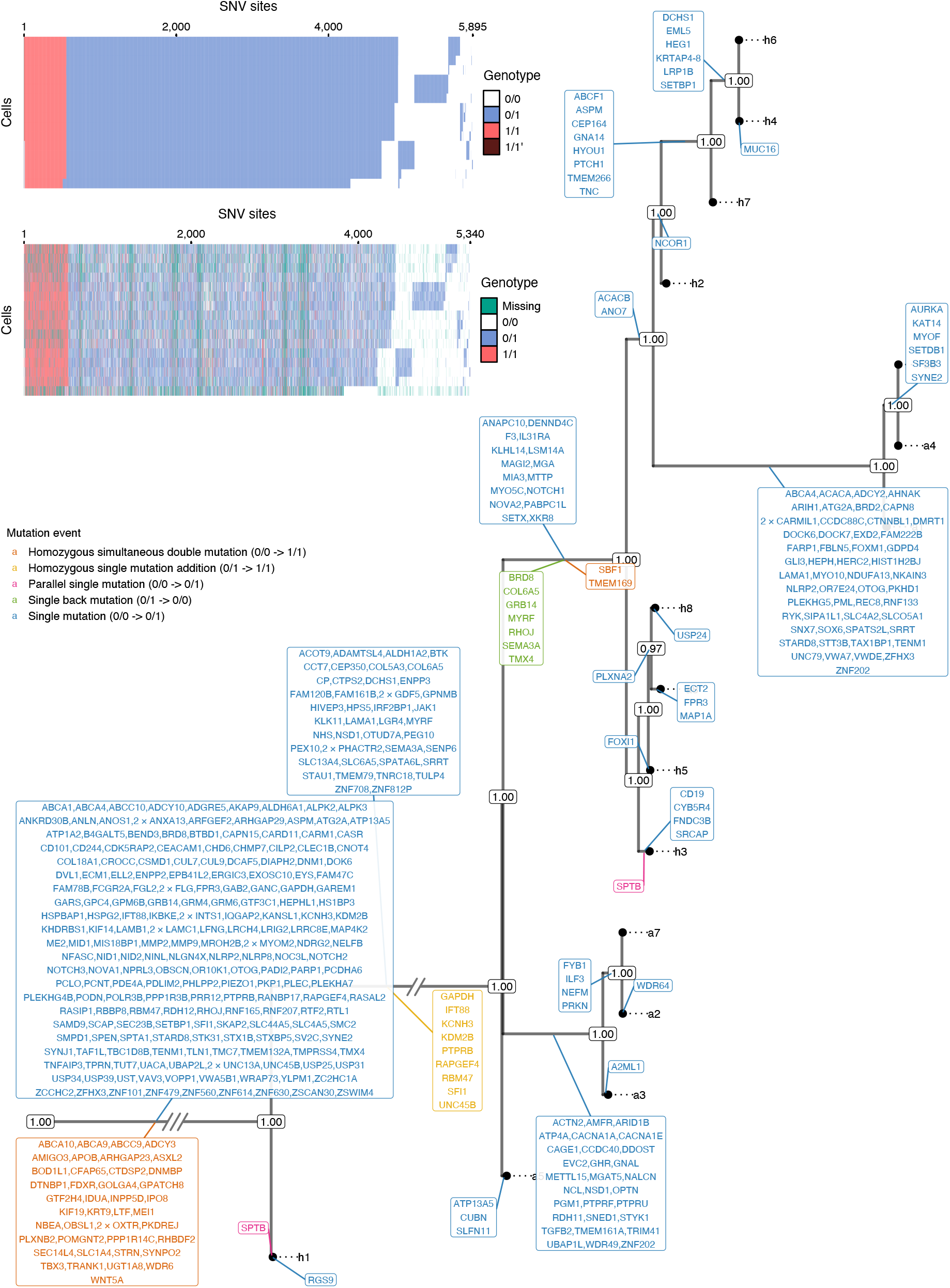
Results of phylogenetic inference and variant calling for TNBC16 [40] dataset. Shown is SIEVE’s maximum clade credibility tree. Two exceptionally long branches are folded with the number of slashes proportional to the branch lengths. Tumour cell names are annotated to the leaves of the tree. The numbers at each node represent the posterior probabilities (threshold *p >* 0.5). At each branch, genes with non-synonymous mutations are depicted in different colours, representing various types of evolutionary events. **a-b**, variant calling heatmap for SIEVE (**a**) and Monovar (**b**). Listed in the legend are the categories of predicted genotypes by each method. Cells in the row are in the same order as that of leaves in the phylogenetic tree.

By mapping non-synonymous mutations to the internal branches, we identified different types of mutation events (Methods), including several violations of the ISA, such as back mutations and parallel mutations. As expected, most of the mutations, including single and double mutant genotypes, resided on the trunk, and some of them occurred in genes which were also reported by the original study [40], such as *TBX3, NOTCH2, NOTCH3* and *SETBP1*. Although SIEVE clustered cells differently from the original study, the high posterior probabilities (Fig. 4) indicate that the tree inferred by SIEVE is more plausible.

SIEVE identified 5,895 SNV sites (Fig. 4a). In contrast to Monovar, SIEVE calls genotypes for all analysed sites, including sites with missing data (Fig. 4b).

### SIEVE inferred a phylogenetic tree and called variants for CRC samples mixed with normal cells

Finally, we applied SIEVE to another scWES dataset [41], which consisted of 48 tumour and normal cells from a patient with CRC (CRC0827 in [41]; referred to as CRC48 below; see Methods). We identified 707 candidate SNV sites as well as 119,486,190 background sites. From the inferred phylogenetic tree (Extended Data Fig. 8 and 9), we inferred two tumour clades matching their anatomical locations (cancer tissue 1 and 2) and one clade for normal cells. Nine cells collected from tumour biopsies were clustered outside the tumour clades, suggesting that these were normal cells within the tumour biopsies. We estimated that the effective sequencing error rate was 8.3 × 10^−4^ and the ADO rate was 0.10.

From the non-synonymous mutations mapped to the branches, we observed unique subclonal mutations, including an established CRC driver mutation, *SYNE1* [39]. We located two parallel single mutations (*CHD3* and *PLD2*), which evolved independently in adenomatous polyps and in tumour cells.

The variant calling results of SIEVE shared a similar but less noisy structure to those of Monovar (Extended Data Fig. 8a,b). We identified 678 SNV sites in total.

## Discussion

Here we present a statistical approach for cell phylogeny inference and variant calling from scDNA-seq data. SIEVE leverages raw read counts to directly reconstruct cell phylogenies and then to reliably call single-cell variants. SIEVE tackles a considerably challenging problem, i.e., the propagation of errors in variant calling to the inference of cell phylogeny, by sharing information between these two tasks. Important characteristics of SIEVE include following the FSA and correction for acquisition bias for tree branch lengths, which prevents from overfitting the phylogenetic model.

Inferring mutation status accurately from highly noisy scDNA-seq data remains a demanding problem. A pivotal strength of SIEVE is its characteristic of using genotypes as a bridge between tree inference and variant calling so that these tasks are united. SIEVE is able to reliably differentiate wildtype, single and double mutant genotypes. The benchmarking shows that SIEVE, regarding variant calling, outperforms methods which employ no cell relationships (Monovar) and which, despite accounting for such information, do not include an instantaneous transition rate matrix and branch lengths (SCIPhI). Regarding tree reconstruction, SIEVE is more robust than SCIPhI, which infers phylogenies following ISA from raw scDNA-seq data. It also outperforms methods that rely on variants called by other approaches as a pre-processing step, thereby likely being misled by wrongly inferred variants (Cellphy and SiFit). The high performance of SIEVE can also be attributed to the fact that it is the only model that performs acquisition bias correction, allowing for more accurate branch lengths, and models the distribution of sequencing coverage and accounting for its overdispersion. Finally, SIEVE is also able to reliably call ADOs given data of adequate coverage quality.

Currently, SIEVE only considers SNVs and assumes a diploid genome. Further improvement could embrace small indels and copy number alterations to improve phylogenetic inference and variant calling, yet care must be taken to differentiate deletions during evolution from ADOs. Additionally, SIEVE only allows at most one ADO for each site and cell. Further extension could expand to locus dropout, which directly results in missing data.

We apply SIEVE to real scDNA-seq datasets harnessed from CRC and TNBC. SIEVE calls far fewer double mutant genotypes and gives more reliable mutation assignment than Monovar does, in line with the simulation results. We also notice that SIEVE identifies double mutant genotypes, which is rare in CRC but frequent in TNBC, indicating the noteworthy role such genotypes play in the evolution of different types of cancer. Future studies could be based on the phylogenetic tree and variants inferred by SIEVE to identify somatic mutations potentially related to the resistance and relapse in the clinical therapy of cancer.

In the real data analysis we utilise the relaxed molecular clock model implemented in BEAST 2. This shows one of the advantages of SIEVE being a package of BEAST 2, and the potential of exploiting the functionality of other BEAST 2 packages in our model. On top of this, SIEVE benefits from the computational efficiency of BEAST 2 solutions, including multi-threaded MCMC.

The SIEVE model successfully exploits raw read counts from scDNA-seq data and jointly infers phylogeny and variants. With the advancement of scDNA-seq technology, we expect the improvement of the coverage quality where the inference of ADO states is reliable. Although we mainly illustrate the application of SIEVE to scDNA-seq data from tumours, it is applicable to studying evolution also in other tissues.

## Methods

### Sample collection

We obtained fresh frozen primary tumour and normal tissues from a single colorectal cancer patient stored at the Galicia Sur Health Research Institute (IISGS) Biobank, member of the Spanish National Biobank Network (Nº B.0000802). This study was approved by a local Ethical and Scientific Committee (CAEI Galicia 2014/015).

### Single-cell isolation, whole-genome amplification and sequencing

We isolated EpCAM+ cells from on normal and three tumoural regions (TP: tumour proximal; TC: tumour central; TD: tumour distal) from the patient with a BD FACSAria III cytometer. We successfully amplified the genomes of 28 cells with Ampli1 (Silicon Biosystems) and built whole-genome sequencing libraries using the KAPA (Kapa Biosystems) library kit. Each library was sequenced at *≈*6x on an Illumina Novaseq 6000 at the Spanish National Center of Genomic Analysis (CNAG-CR; https://www.cnag.crg.eu/). We called this dataset CRC28.

### Data preprocessing

For the public TNBC16 [40] and CRC48 [41] datasets, we downloaded the raw sequencing reads from the SRA database in FASTQ format. For the three datasets (CRC28, TNBC16 and CRC48) We trimmed the Illumina adapter sequences using cutadapt (version 1.18) and mapped reads to the 1000G Reference Genome hs37d5 using BWA MEM (version 0.7.17). After de-duplication with Picard (version 2.18.14), we used GATK (version 3.7.0) for local realignment based on indel calls from the 1000G Phase 1 and the Mills and 1000G gold standard. Subsequently, we recalibrated the base scores using GATK (version 4.0.10) with polymorphisms from dbSNP (build 138) and indels from the 1000G Phase 1. Exact commands used to run the tools are featured in Supplementary Note.

### SIEVE model

SIEVE is a statistical approach which combines a statistical phylogenetic model with a probabilistic model of raw read counts. We implement SIEVE under BEAST 2 [34], a popular Bayesian phylogenetic framework that uses Markov Chain Monte Carlo (MCMC) for the estimation of phylogenetic trees and model parameters.

### Input data

SIEVE takes as input raw read counts of all four nucleotides at candidate SNV sites (Fig. 1a). Specifically, for cell *j ∈ {*1, …, *J}* at candidate SNV site *i ∈ {*1, …, *I}*, the input data to SIEVE is in the form of 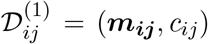, where ***m***_***ij***_ = *{m*_*ijk*_ | *k* = 1, 2, 3*}* corresponds to the read counts of three alternative nucleotides with values in descending order and *c*_*ij*_ to the sequencing coverage for cell *j* and site *i*.

Candidate SNV sites are defined as statistically significant SNVs. They are referred to as ‘candidate’ since this significance could sometimes be a false discovery due to technical errors in scDNA-seq. To identify the candidate SNV sites we developed a tool named DataFilter that employs a strategy similar to SCIPhI [15]. Specifically, a likelihood ratio test is conducted for SNV detection, but with a modification enabling to capture sites containing double mutant genotypes.

For scWGS and scWES datasets, raw read counts from *I*^′^ background sites are denoted *𝒟*^(2)^. The number of background sites is used to correct acquisition bias (see Section SIEVE likelihood). For datasets lacking background information (for instance, from targeted sequencing), SIEVE accepts a user-specified number of background sites only for acquisition bias correction.

### Statistical phylogenetic model

The statistical phylogenetic model behind SIEVE includes an instantaneous transition rate matrix, which is defined by a continuous-time homogeneous Markov chain. We consider four possible genotypes *G* = {0/0, 0/1, 1/1, 1/1^′^*}*, where 0, 1, and 1^′^ are used to denote the reference nucleotide, an alternative nucleotide, and a second alternative nucleotide which is different from that denoted by 1, respectively. The fundamental evolutionary events we consider are single mutations and single back mutations. The former happen when 0 mutates to 1, or 1 and 1^′^ mutate to each other, while the latter occur when 1 or 1^′^ mutates to 0. Hence, genotypes 0/0 and 0/1 represent wildtype and single mutant genotypes, respectively, whereas genotype 1/1 and 1/1^′^ represent double mutant genotypes. We intentionally use the non-standard nomenclature of single and double mutants to discern important evolutionary events. In contrast, calling both 0/1 and 1/1’ a heterozygous mutation genotype would be more standard and correct, but would not differentiate between the genotype that has only a single allele changed with respect to the reference (0/1) from the genotype that has two alleles changed (1/1’). We only consider unphased genotypes, so we do not differentiate between 0/1 and 1/0 or between 1/1^′^ and 1^′^/1.

The joint conditional probability of all cells at SNV site *i* having genotype *g*_*ij*_ *∈ G, j* = 1, …, *J* is determined according to the statistical phylogenetic model by

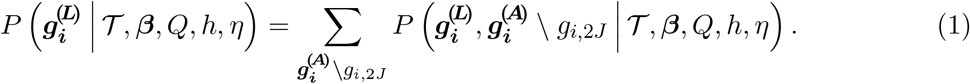

In Eq. (1), ***β*** represents the branch lengths measured by the expected number of somatic mutations per site and *Q* is the instantaneous transition rate matrix of the Markov chain. *𝒯* is the rooted binary tree topology, representing the genealogical relations among cells. We specifically require the root of *𝒯* to have only one child, representing the most recent common ancestor (MRCA) of all cells. The branch between the root and the MRCA is the trunk of the cell phylogeny. The trunk is one of novelties of our approach, introduced to represent the accumulation of clonal mutations (shared among all cells) in the initial phase of tumour progression. Therefore, with *J* existing cells, labelled by {1, …, *J*}, as leaves, *𝒯* has *J* internal hidden ancestor nodes, labelled by *{J* + 1, …, 2*J*}, and 2*J −* 1 branches, whose lengths are kept in ***β***. The trunk is essential for *𝒯* to assure that the root, labelled by 2*J*, represents a normal ancestor cell even if the data only contains tumour cells. Hence the genotype of the root for SNV site *i*, denoted *g*_*i*,2*J*_, is fixed to 0/0. 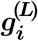 represents the genotypes of *J* cells as leaves of *𝒯*, while 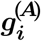 is the genotypes of all ancestor cells as internal nodes of *𝒯*. Note that we marginalise the genotypes of the ancestor nodes except for the root. We also consider among-site substitution rate variation following a discrete Gamma distribution with mean equal 1, parameterised by the number of rate categories *h* and shape *η* [42]. *𝒯*, ***β***, *η* in Eq. (1) are hidden variables, estimated using MCMC (see Section Posterior and MCMC), whereas *h* is a hyperparameter that is fixed (4 by default). Note that variant calling effectively corresponds to the determination of the values of the variables 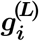.

In the transition rate matrix *Q* (Fig. 1c), each entry denotes a rate from one genotype to another during an infinitesimal time interval Δ*t*. Note that at most one change is allowed to occur in Δ*t*. For instance, the transition of 0/0 moving to 1/1 during Δ*t* is impossible as two single somatic mutations are required; thus, the corresponding transition rate is 0. The transition rate from genotype 0/0 to 0/1 represents the somatic mutation rate and is set to 1. The back mutation rate is measured relatively to the somatic mutation rate and therefore is ^1^/_3_.

With the genotype state space *G* defined, for a given branch length *β*, the underlying fourby-four transition probability matrix *R*(*β*) of the Markov chain is represented using matrix exponentiation of the product of *Q* and *β* as *R*(*β*) = exp(*Qβ*) [27].

### Model of raw read counts

The probability of observing the input data *𝒟*_*ij*_ for cell *j* at site *i* is factorized as

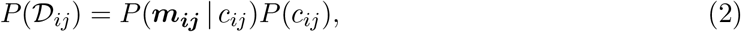

where the first component is the model of nucleotide read counts and the second the model of sequencing coverage.

### Model of sequencing coverage

After single-cell whole-genome amplification (scWGA) some genomic regions are more represented than others. After scDNA-seq, this results in an uneven coverage along the genome, much more than in the case of bulk sequencing. Here, to model the sequencing coverage *c* in the presence of overdispersion, we employ a negative binomial distribution.

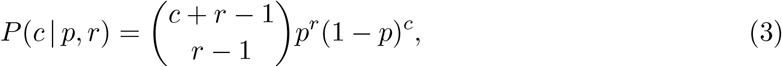

with parameters *p* and *r*. We reparameterise the distribution with 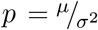 and 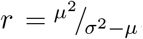, where *μ* and *σ*^2^ are the mean and the variance of the distribution of the sequencing coverage *c*, respectively.

Theoretically, each cell *j* at site *i* has its specific *μ*_*ij*_ and 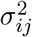 parameters, which, however, are impossible to be estimated freely. Hence, we make additional assumptions and pool the data for better estimates, adapting the approach of [43]. We assume that *μ*_*ij*_ and 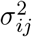 have the following forms, respectively:

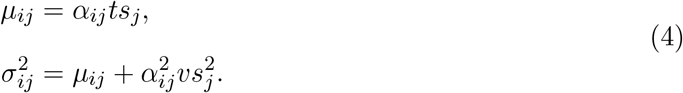

In Eq. (4), *t* is the mean of allelic coverage (the expected coverage per allele) and *v* is the variance of allelic coverage. We estimate *t* and *v* with MCMC (see Section Posterior and MCMC). *α*_*ij*_ *∈ {*1, 2*}* is a hidden random variable denoting the number of sequenced alleles for cell *j* at site *i*. According to the statistical phylogenetic model, both alleles are expected to be sequenced. However, due to the frequent occurrence of allelic dropout (ADO) during scWGA, there are cases where only one allele is amplified and therefore *α*_*ij*_ is 1. Eq. (4) reflects the fact that the expected sequencing coverage and its raw variance are proportional to the number of sequenced alleles. Note that inferring the hidden variable *α*_*ij*_ corresponds to identifying occurrences of ADO events, and hence the ability of SIEVE to perform ADO calling. We denote the prior distribution of *α*_*ij*_

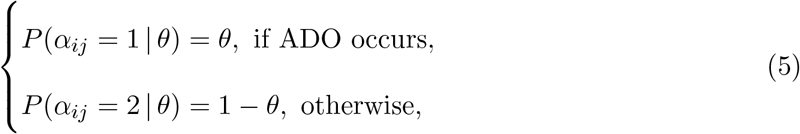

where *θ* is a parameter corresponding to the the probability of ADO occurs, i.e., the ADO rate, which is estimated using MCMC.

In Eq. (4), *s*_*j*_ is the size factor of cell *j* which makes sequencing coverage from different cells comparable and is estimated directly from the sequencing coverage using

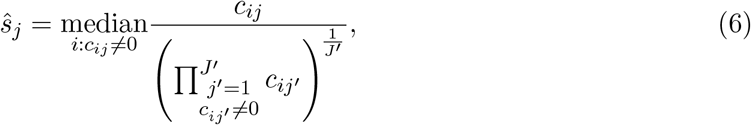

where *J*^′^ is the number of cells with non-zero coverage at a site. By taking into account only the non-zero values, the estimate *ŝ*_*j*_ is not affected by the missing data, which is prevalent in scDNA-seq.

### Model of nucleotide read counts

We denote the genotype affected by ADO 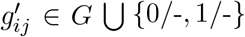, where 0/-and 1/-are the results of ADO occurring to *g*_*ij*_. For instance, 0/-is caused either by 0 dropped out from 0/0 or by 1 dropped out from 0/1. Then the probability of 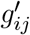 is denoted by

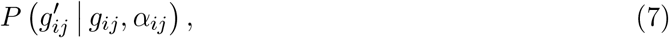

which is defined at length in Table 1.

We model the read counts of three alternative nucleotides ***m***_***ij***_ given the sequencing coverage *c*_*ij*_ with a Dirichlet-multinomial distribution as

**Table 1:** Definition of the distribution of 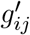 conditional on *g*_*ij*_ and *α*_*ij*_.

| $g'_{ij}$ | $g_{ij}$ | $\alpha_{ij}$ | $P(g'_{ij} \vert g_{ij}, \alpha_{ij})$ |
| --- | --- | --- | --- |
| 0/0 | 0/0 | 2 | 1 |
| 0/- | 0/0 | 1 | 1 |
| 0/1 | 0/1 | 2 | 1 |
| 1/1 | 1/1 | 2 | 1 |
| 1/- | 1/1 | 1 | 1 |
| 1/1' | 1/1' | 2 | 1 |
| 1/- | 1/1' | 1 | 1 |
| 0/- | 0/1 | 1 | $\frac{1}{2}$ |
| 1/- | 0/1 | 1 | $\frac{1}{2}$ |
| Others |  |  | 0 |

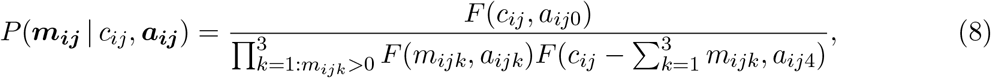

with parameters ***a***_***ij***_ = *{a*_*ijk*_ | *k* = 1, …, 4*}* and 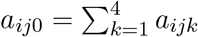. *F* is a function in the form of

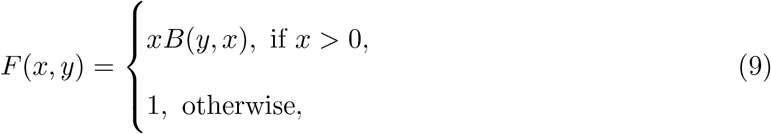

where *B* is the beta function. Note that 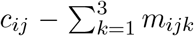 is the read count of the reference nucleotide.

To improve the interpretation of Eq. (8), we reparameterise it with ***a***_***ij***_ = *w*_*ij*_***f***_***ij***_, where ***f***_***ij***_ = *{f*_*ijk*_ | *k* = 1, …, 4*}*, 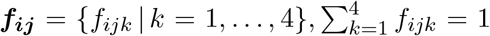 is a vector of expected frequencies of each nucleotide and *w*_*ij*_ represents overdispersion. ***f***_***ij***_ are categorical hidden variables dependent on 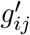:

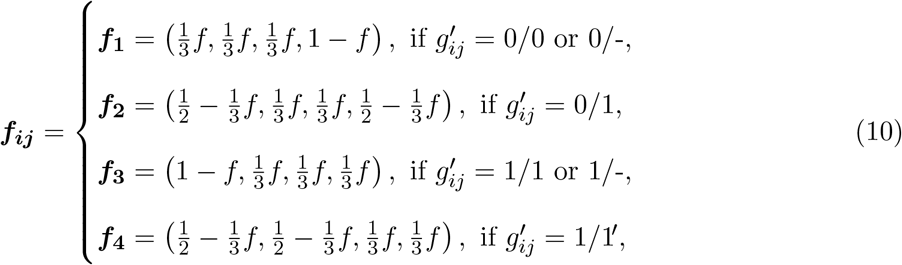

where *f* is the expected frequency of nucleotides whose existence is solely due to technical errors during sequencing. To be specific, *f* is defined as the effective sequencing error rate including amplification (where a nucleotide is wrongly amplified into another one during scWGA) and sequencing errors.

*w*_*ij*_ is also a categorical hidden variable dependent on 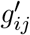:

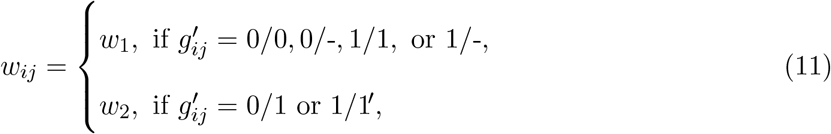

where *w*_1_ is wild type overdispersion and *w*_2_ is alternative overdispersion.

By plugging in Eqs. (10) and (11), Eq. (8) is equivalently represented with

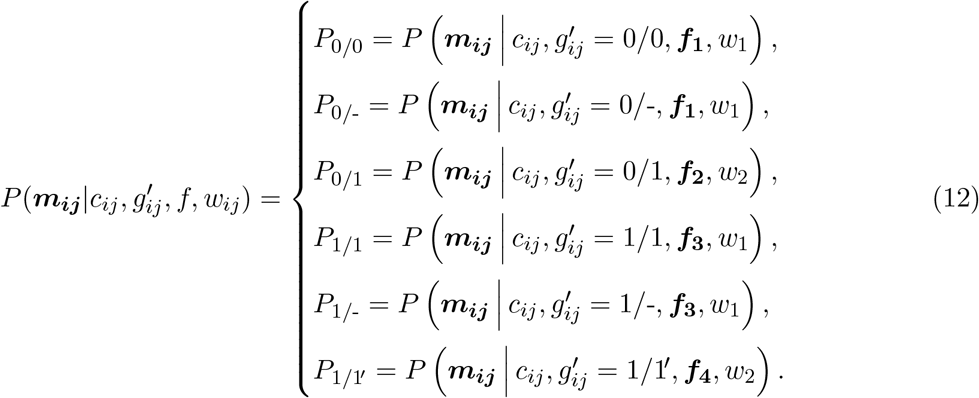

Note that *P*_0/0_ and *P*_0/-_ share the same ***f*** and *w*_1_, showing that the model of nucleotide read counts is not enough to discriminate 0/0 from 0/-, and so do *P*_1/1_ and *P*_1/-_. In such cases, incorporating the model of sequencing coverage helps resolve the entanglement.

To understand Eq. (12), first take *P*_0/0_ as an example. Theoretically, no alternative nucleotides are supposed to exist if no technical errors occur. Thus, any observations of any alternative nucleotides can only result from technical errors, and the expected frequency of the reference nucleotide is accordingly adjusted to 1−*f*. For another example *P*_0/1_, say the reference nucleotide is A and the alternative nucleotide is C, and both their read count frequencies are supposed to be ^1^/_2_ if no technical errors occur. For the other two alternative nucleotides, G and T, their observations could only result from technical errors, and both their frequencies are ^*f*^/_3_. Moreover, either A or C may be sequenced as a different nucleotide (each with probability 1/2). In the former case, the frequency of A decreases by ^*f*^/_2_. In the latter case, if C is sequenced as A (with probability ^*f*^/_3_) the frequency of A increases by ^1^/_2_ × ^*f*^/_3_. Overall, the frequency of A decreases by ^*f*^/_3_, resulting in ^1^/_2_ − ^*f*^/_3_.

*f, w*_1_ and *w*_2_ in Eq. (12) are estimated with MCMC.

### SIEVE likelihood

We denote the conditional variables in Eq. (1) as Θ = *{𝒯*, ***β***, *Q, h, η}* and those in the model of raw read counts as Φ = *{t, v, θ, f, w*_1_, *w*_2_*}*. Given the input data *𝒟*^(1)^ and *𝒟*^(2)^, the log-likelihood of the SIEVE model is

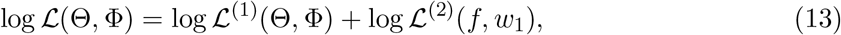

where *ℒ*^(1)^ is the tree likelihood corrected for acquisition bias computed from candidate SNV sites in *𝒟*^(1)^, while *ℒ*^(2)^ is the likelihood computed from background sites in *𝒟*^(2)^, referred to as the background likelihood. Eq. (13) does not contain *g*_*ij*_, 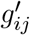, *α*_*ij*_ since they are marginalised out (see below).

Since we only use data from SNV sites to compute the tree likelihood, the tree branch lengths ***β*** are prone to be overestimated [29, 30]. The overestimation of ***β*** due to only using data from SNV sites is called acquisition bias, which is corrected in SIEVE according to [44]:

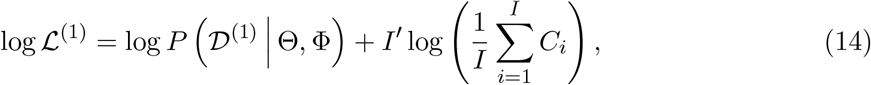

where the first component is the uncorrected tree log-likelihood for SNV sites, and *C*_*i*_ in the second component is the likelihood of SNV site *i* being invariant (see below). The regularization term 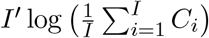 renders SIEVE in favor of trees with short branch lengths where *ℒ*^(1)^ is large due to the increasing averaged *C*.

To compute the uncorrected tree log-likelihood, we marginalise out *α*_*ij*_ and 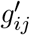 :

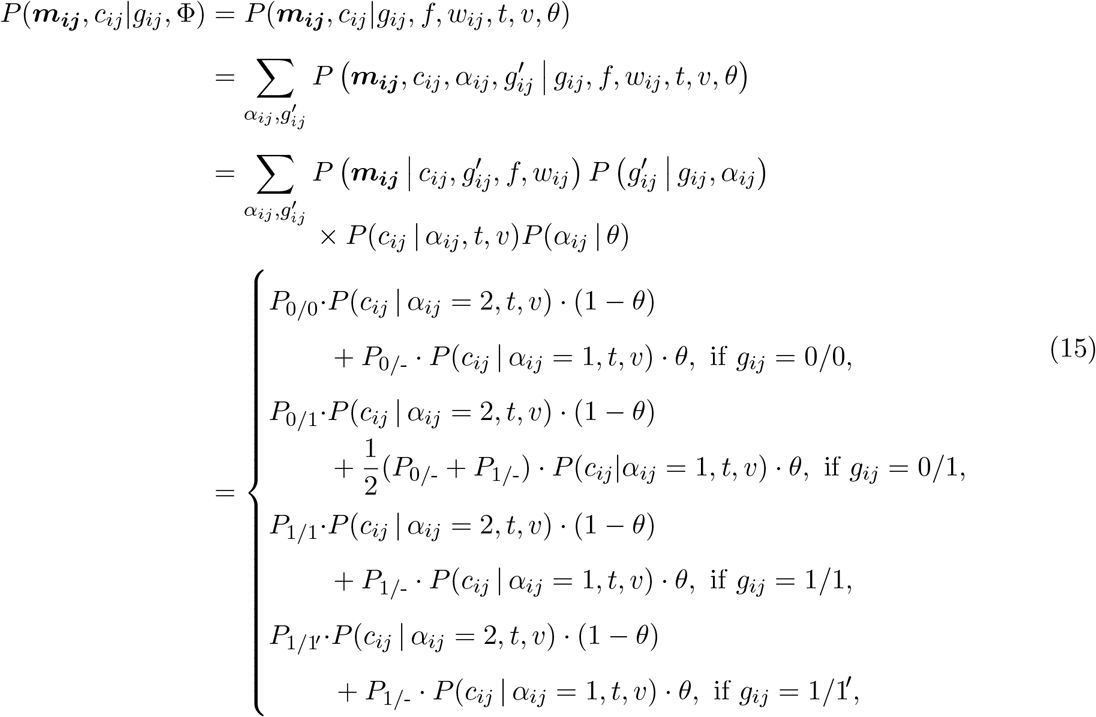

where *P*_0/0_, *P*_0/-_, *P*_0/1_, *P*_1/1_, *P*_1/-_, *P*_1/1_^′^ are defined in Eq. (12) and 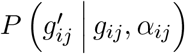 is defined in Eq. (7). In the second line of Eq. (15), the probability is factorised out according to Fig. 1b.

To compute log *P*(*𝒟*^(1)^ Θ, Φ) in Eq. (14), we assume that the SNV sites evolve independently and identically. By plugging Eqs. (1) and (15), log *P*(*𝒟*^(1)^ Θ, Φ) is denoted by

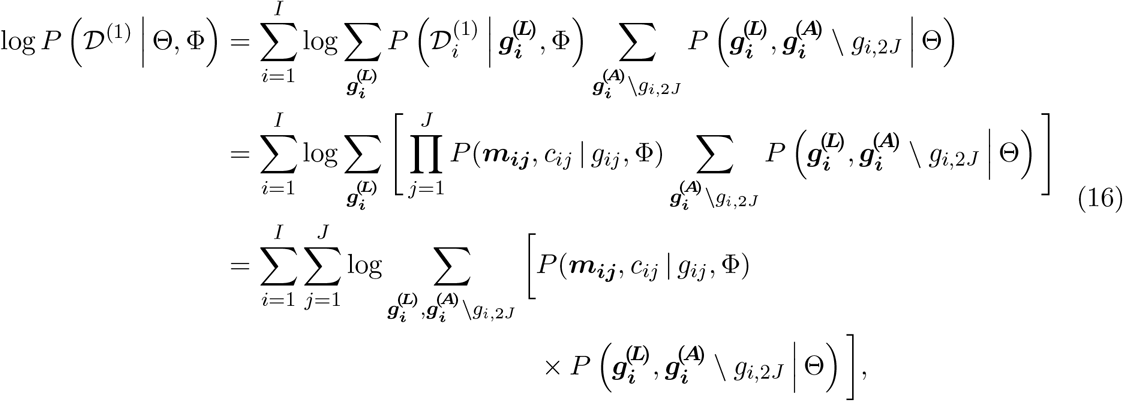

which is efficiently computed out by Felsenstein’s pruning algorithm [45], with the extension of the model of raw read counts applied on leaves. Specifically, the Fenselstein’s pruning algorithm is applied to an extended tree *𝒯*, where additional leaf nodes corresponding to the data are attached at the bottom of *𝒯* : for each node corresponding to genotype *g*_*ij*_ there is a leaf node added, corresponding to data (***m***_***ij***_, *c*_*ij*_), and the transition probability between the genotype node and the leaf is given by Eq. (15). For *I* candidate SNV sites, *J* cells and *K* genotype in *G* (for SIEVE *K* = 4), the time complexity of Felsenstein’s pruning algorithm is *𝒪*(*IJK*^2^).

*C*_*i*_ in Eq. (14) is determined similarly to Eq. (16) by computing the joint probability of observing the data 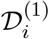 and 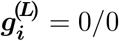:

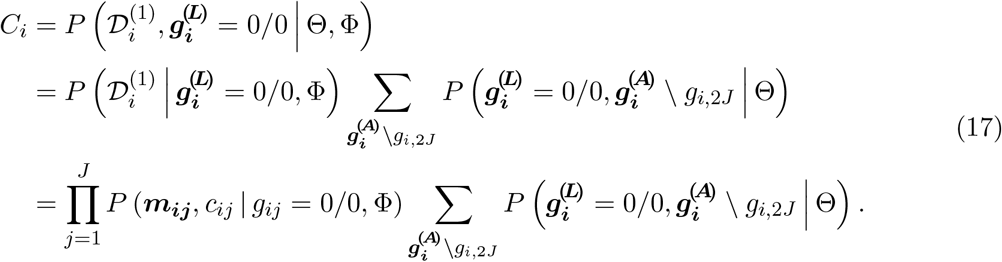

Formally, to compute the background likelihood, we should account for the fact that the background sites, similarly to the variant sites, also evolve under the phylogenetic model and involve similar computations as above. This, however, would result in a large additional computational burden due to the large number of background sites compared to the variant sites. Thus, to estimate the background log-likelihood efficiently, we make several simplifications and compute it only approximately. First, we assume that across *I*^′^ background sites each cell has the same genotype 0/0 and both alleles are covered. We further ignore the model of sequencing coverage and the tree log-likelihood in the computations. As a result, by employing an alternative expression of Dirichlet-multinomial distribution log *ℒ*^(2)^ is efficiently obtained as

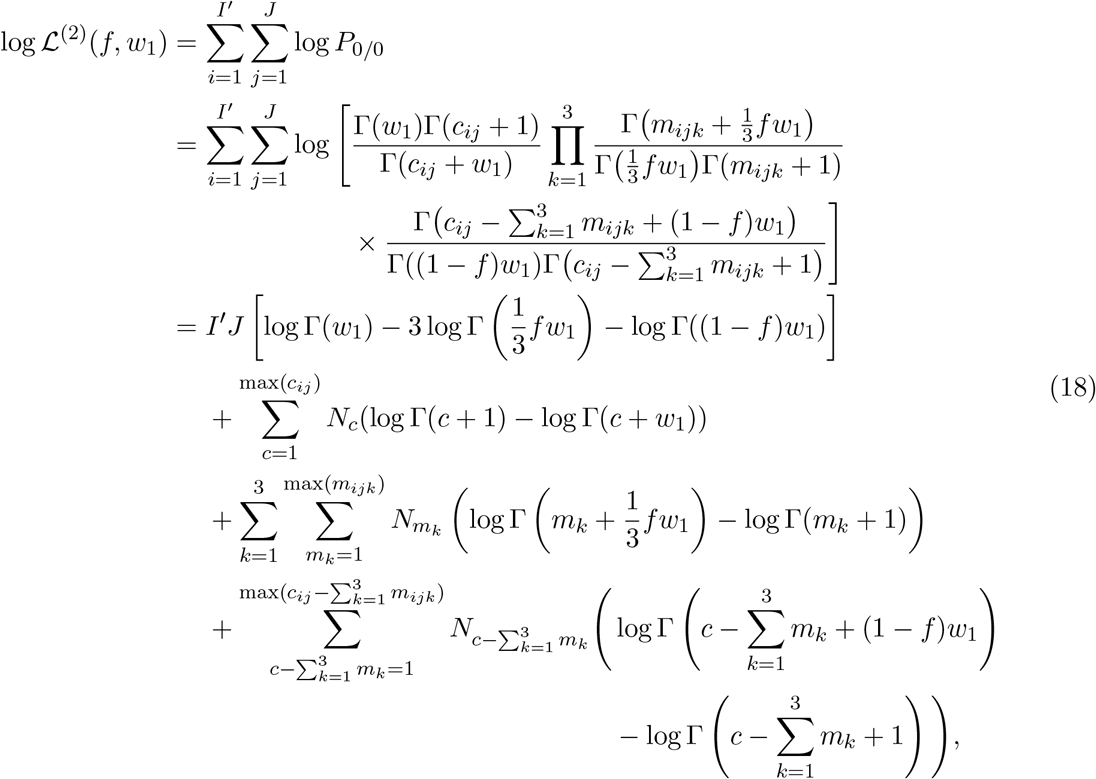

where *P*_0/0_ is defined in Eq. (12). *N*_*c*_, 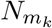 for *k* = 1, 2, 3, and 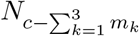 represent, across *I*^′^ background sites and *J* cells, the unique occurrences of sequencing coverage *c*, of alternative nucleotide read counts *m*_1_, *m*_2_, *m*_3_, and of reference nucleotide read counts 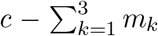, respectively. In Eq. (18), some items, namely log Γ(*c* + 1), − log Γ(*m*_*k*_ + 1) for *k* = 1, 2, 3, and 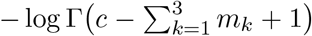, only depends on the data, which remain constants during MCMC. Therefore, they are ignored in the computation of background likelihood. It is clear that the background likelihood helps estimate *f* and *w*_1_.

The time complexity of Eq. (18) is *𝒪*(*c*) with *c* being the number of unique values of sequencing coverage across all cells and background sites. Since *IJK*^2^ is usually much larger than *c*, the overall time complexity of model likelihood is *𝒪*(*IJK*^2^).

### Priors

To define priors for model parameters and for the tree coalescent, we employ the prior distributions defined in BEAST 2. We impose on *𝒯* and ***β*** in Eq. (1) a prior distribution following the Kingman coalescent process with an exponentially growing population. The tree prior is parameterised by scaled population size *M* and exponential growth rate *q*, and is denoted by

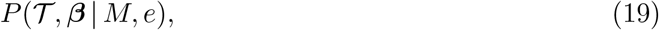

whose analytical form is defined in [46]. *M* and *e* are hidden random variables and are estimated using MCMC. Note that, by default, *M* represents the number of time units, e.g., the number of years, and the mutation rate is measured by the number of mutations per time unit per site. Their product results in the unit of branch length, i.e., the number of mutations per site. Since scDNA-seq data usually does not contain temporal information as a result of collecting samples at the same time, it is impossible to differentiate *M* from the mutation rate. However, if the mutation rate is known, one could alternatively estimate a time-calibrated cell phylogeny.

As prior distributions, we assign to *M*

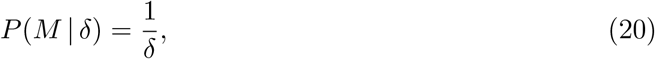

where *δ* is the current proposed value of *M*. Note that this is supposed to be normalised to define a proper probability distribution, but this form is sufficient to define a proper posterior (see Section Posterior and MCMC).

For *e* we choose

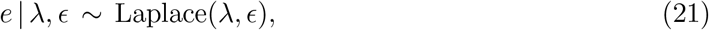

where we choose mean *λ* = 10^−3^ and scale *ϵ* = 30.7 (default in the BEAST 2 software). We choose an exponential distribution as the prior for *η* in Eq. (1):

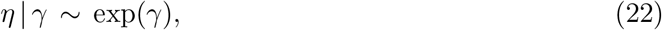

where *γ* = 1.

For the model of sequencing coverage described in Eqs. (3) and (4), we set the prior for *t* within a large range of values with

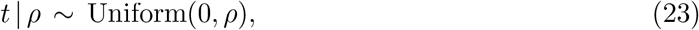

where *ρ* = 1000, and the prior for *v* with

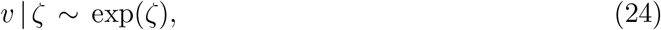

where *ζ* = 25. In terms of *θ* in Eq. (5), it also has a uniform prior:

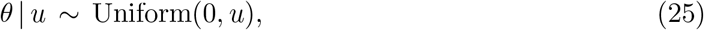

where *u* = 1.

For the model of nucleotide read counts described in Eqs. (10) to (12), we choose an exponential prior for *f* :

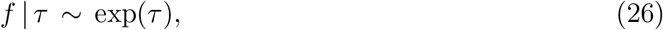

where *τ* = 0.025, and a log normal prior for both *w*_1_ and *w*_2_:

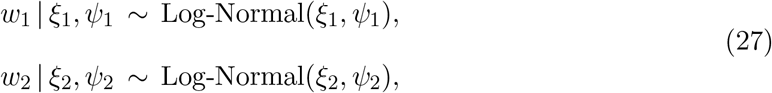

where we choose for *w*_1_ the mean *ξ*_1_ = 3.9 and the standard deviation *ψ*_1_ = 1.5, and for *w*_2_ the mean *ξ*_2_ = 0.9 and the standard deviation *ψ*_2_ = 1.7. These specific values reflect our belief that *w*_1_ is greater than *w*_2_, while both distributions cover a large range of possible values for *w*_1_ and *w*_2_.

### Posterior and MCMC

With the model likelihood and priors defined, the posterior distribution of the unknown parameters is

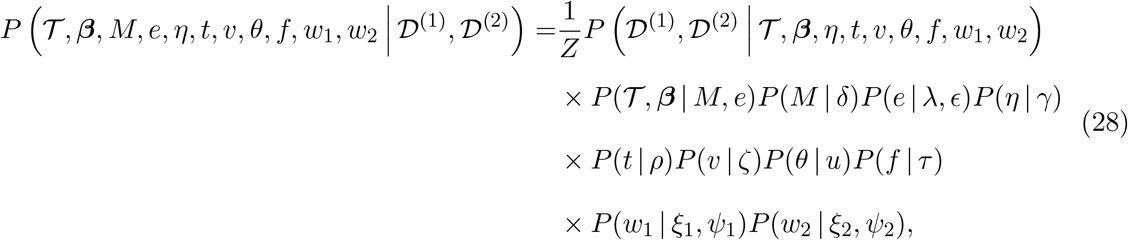

where *Z* is a normalisation constant, representing the probability of the observed data.

Since the posterior distribution does not have a closed-form analytical formula, we employ the MCMC algorithm with Metropolis-Hastings kernel to sample from the posterior distribution in Eq. (28). Given the current state of the parameters *q*, we propose a new state *q*^*\**^ according to proposal distributions *P* (*q*^*\**^|*q*) that assure the reversibility and ergodicity of the Markov chain. With one parameter changed a time, *q*^*\**^ is accepted with probability

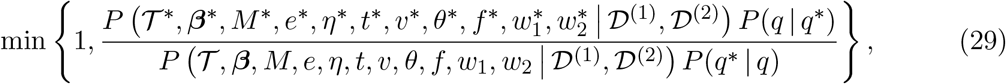

where the normalisation constant *Z* cancels out after plugging in Eq. (28).

For sampling the structure of the cell phylogeny, we take advantage of proposal distributions implemented in the BEAST 2 software [46] and modify them to make sure they are compatible with our tree topology, so that the sampled trees are binary and contain a trunk. Specifically, the tree branch lengths are changed by scaling the heights of the internal nodes. For tree topological exploration, we use the Wilson-Balding move to perform subtree pruning and regrafting. Specifically, a random node and half of its subtree is pruned and reattached to a random branch not belonging to the moved subtree. A subtree-slide move is also used, where a random node and half of its subtree slides either upwards or downwards along branches and cross at least one node. Both those two moves include changes to the lengths of some branches. The final type of move swaps two randomly selected subtrees.

For sampling unknown parameters, we perform either scaling operations or random Gaussian walks.

SIEVE runs with a two-stage sampling strategy. In the first stage the acquisition bias correction is switched off and all parameters are explored, while in the second stage the acquisition bias correction is turned on and parameters not affecting branch lengths are fixed with their estimates from the previous stage. This two-stage strategy proved to yield more accurate parameter and tree estimates than a strategy where both parameters and tree would be explored at once, with the acquisition bias correction enabled. Additionally, the initial tree in the second stage is set to the tree summarised from the first stage.

### Variant calling, ADO calling and maximum likelihood gene annotation

During the sampling process 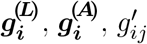 and *α*_*ij*_ (Eqs. (1), (15) and (16)) are hidden variables that are marginalised out. Therefore, to obtain estimates of these hidden variables, we infer their maximum likelihood configuration with the max-sum algorithm [47], using the maximum clade credibility tree [48] and parameters estimated from the MCMC posterior samples.

To be specific, by determining the maximum likelihood genotypes of the leaves 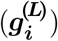, we are able to call variants. By inferring the maximum likelihood 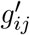 and *α*_*ij*_, the ADO state is determined. Moreover, by computing the maximum likelihood genotypes of the internal nodes 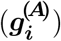, SIEVE maps mutations to specific tree branches. Mutation events are classified into different categories (see Table 2).

**Table 2:** Twelve types of mutation categories that SIEVE is able to identify.

| Genotype transition | Mutation event |
| --- | --- |
| 0/0 → 0/1 | Single mutation |
| 0/0 → 1/1 | Homozygous simultaneous double mutation |
| 0/0 → 1/1' | Heterozygous simultaneous double mutation |
| 0/1 → 0/0 | Single back mutation |
| 1/1 → 0/1 | Single back mutation |
| 1/1' → 0/1 | Single back mutation |
| 0/1 → 1/1 | Homozygous single mutation addition |
| 0/1 → 1/1' | Heterozygous single mutation addition |
| 1/1 → 0/0 | Double back mutation |
| 1/1' → 0/0 | Double back mutation |
| 1/1' → 1/1 | Homozygous substitute single mutation |
| 1/1 → 1/1' | Heterozygous substitute single mutation |

### scDNA-seq data simulator

In order to benchmark the performance of SIEVE against those of other published methods, we simulated scDNA-seq data by modifying CellCoal [49] (commit 594e063). In contrast to CellCoal, the sequencing coverage is generated according to Eqs. (3) to (6). Given the sequencing coverage, read counts are simulated with a Multinomial distribution including errors. Input configuration follows the one described for CellCoal [49].

The simulator mimics both the biological evolution and the sequencing process. We first generated a binary genealogical cell lineage tree following the coalescent process assuming a strict molecular clock and created a reference genome where each site was initialised by the reference genotype with one of the four nucleotides. With a specific mutation rate, each site was evolved independently along the tree according to a rate matrix which contains ten diploid genotypes encoded with nucleotide pairs (Supplementary Table 2). The rate matrix allows mutations and back mutations, where the probability of the latter is ^1^/_3_ of the former. All simulated sites for which at least one cell has a non-reference genotype are considered as true SNV sites. Next, we added at most one ADO to cell *j* at site *i* according to the ADO rate. If ADO happens, the number of sequenced alleles *α*_*ij*_ drops from two to one. We recorded the true ADO states across cells for the SNV sites. Size factors for cells in Eq. (4) were sampled from a normal distribution (mean = 1.2, variance = 0.2). Using the negative binomial distribution, we simulated the sequencing coverage with given *t* and *v*. Based on the ADO-affected genotype and sequencing coverage, the read count for each nucleotide was simulated using a Multinomial distribution with a given amplification error rate and sequencing error rate.

### Simulation design

We designed simulations to compare multiple methods in different aspects. We assumed that the tumour cell samples belonged to an exponentially growing population (growth rate = 10^−4^) with an effective population size of 10^4^. The number of tumour cells was chosen to be either 40 or 100. We selected three mutation rates: 10^−6^, 8 × 10^−6^, and 3 × 10^−5^. For different mutation rates, different total number of sites were chosen to result in around 1000 SNV sites for 100 cells (1.3 × 10^5^ sites for 10^−6^, 2 × 10^4^ sites for 8 × 10^−6^, and 6.5 × 10^3^ sites for 3 × 10^−5^), as well as between 250 to 1000 SNV sites for 40 cells (8 × 10^4^ sites for 10^−6^, 2 × 10^4^ sites for 8 × 10^−6^, and 5 × 10^3^ sites for 3 × 10^−5^). Additionally, we varied *t* and *v* in Eqs. (3) and (4) to simulate different coverage qualities. For high quality data, we chose high mean (*t* = 20) and low variance (*v* = 2) of allelic coverage. For medium quality data, we chose high mean (*t* = 20) and medium variance (*v* = 10). For low quality data, we chose low mean (*t* = 5) and high variance (*v* = 20), which was specifically created to mimic the CRC28 dataset.

Other important parameters in the simulation were fixed as follows: in Eq. (5) *θ* = 0.163, in Eq. (12) *w*_1_ = 100 and *w*_2_ = 2.5, and both amplification error rate and sequencing error rate were 10^−3^, which resulted in the effective sequencing error rate *f ≈* 2 × 10^−3^ in Eq. (12).

We designed in total 18 simulation scenarios, each repeated 20 times. The benchmarking framework was built using Snakemake [50].

### Measurement of cell phylogeny accuracy and quality of variant calling

To assess the accuracy of the cell phylogeny reconstruction considering branch lengths, we computed the rooted BS distance from the inferred tree to the true tree [35]. For any two trees, this difference is computed as:

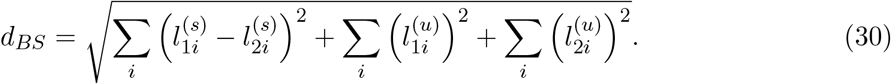

where 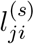 represents the length of a branch shared by both trees, and 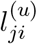 represents the length of a branch *i* that is unique for tree *j*.

To assess the accuracy of the cell phylogeny reconstruction ignoring branch lengths we used the normalised RF distance [36]:

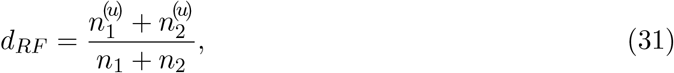

where *n*_*j*_ denotes the total number of branches in tree *j*, while 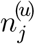 represents the number branches exclusive of tree *j*.

Thus, rooted BS distance and normalised RF distance values equal to 0 indicate a perfect tree reconstruction. For SIEVE and SiFit, we compute both normalised RF distance and rooted BS distance in the rooted tree mode. For CellPhy, we compute these metrics in the unrooted tree mode as it infers an unrooted tree from data only containing tumour cells. Since SCIPhI reports a rooted tree without branch lengths, we can only compute the normalised RF distance. Rooted BS distance and normalised RF distance values were computed using the R package phangorn [51].

To evaluate the variant calling and ADO calling results, we computed precision, recall, F1 score and false positive rate (FPR). For variant calling, we separately compared the performance in calling the single mutant genotype and double mutant genotypes. In particular, when we evaluated the accuracy of single mutant genotype calling, any identification of double mutant genotypes whose true genotype is single mutant genotype was counted as a false negative. Moreover, we analysed two different types of false positives in single mutant genotype calling. The first type corresponds to single mutation calls for sites where the true genotype is a wildtype genotype. The second type are single mutant calls for sites where the true genotype is a double mutant.

For SIEVE and Monovar, we computed the recall, precision, F1 score, and FPR for single mutant genotype calling and double mutant genotype calling. For SCIPhI, we only computed metrics for single mutant genotype calling as it does not call double mutant genotypes. Moreover, we evaluated the accuracy of calling ADO states only for SIEVE, as it is the only method that is able to call them.

### Configurations of methods

For Monovar (commit 68fbb68), we used the true values of *θ* and *f* as priors for false negative rate and false positive rate and default values for other options.

For SCIPhI (commit 34975f7), we ran it with default options and 5 × 10^5^ iterations.

To run CellPhy (commit 832f6c2) and SiFit (commit 9dc3774), we fed the required data with variants called by Monovar. For CellPhy, we piped the data in VCF format and initialised the tree search with three parsimonious trees. We instructed the tool to use a built-in rate matrix with ten genotypes (GT10), a stationary nucleotide frequency distribution learned from the data (FO), an error model applied to the leaves (E), and the Gamma model of site-wise substitution rate variation (G). For SiFit, we fed the input data as a ternary matrix and used the true values of *θ* and *f* as the prior for false negative rate and the estimated false positive rate, respectively. We ran it with 2 × 10^5^ iterations.

On the simulated data, we ran SIEVE with a strict molecular clock model for 2 × 10^6^ and 1.5×10^6^ iterations for the first and the second sampling stage, respectively. On the real datasets, we used a log-normal relaxed molecular clock model to take into consideration branch-wise substitution rate variation. To achieve better mixed Markov chains, we employed a optimized relaxed clock model in [37] instead of the default one in BEAST 2.

Since more parameters are added when using the relaxed molecular clock model, we ran the analysis with 3 × 10^6^ iterations for the first stage and 2.5 × 10^6^ iterations for the second, respectively. Note that the parameters introduced by the relaxed molecular clock model are also explored in the second sampling stage. The SNVs were then annotated using Annovar (version 2020 Jun. 08) [52]. In the main text, the tree was plotted using ggtree [53] and the genotype heatmap was plotted using ComplexHeatmap [54].

## Data availability

Raw single-cell whole-genome sequencing data from CRC28 have been deposited in the Sequence Read Archive (SRA, https://www.ncbi.nlm.nih.gov/sra) database under the accession code XXXXX. We have additionally analysed two published single-cell datasets ([40, 41]). Raw sequencing data for these datasets are available from the SRA database under accession codes SRA053195 (TNBC16) and SRP067815 (CRC48).

## Code availability

SIEVE is implemented in Java and is accessible at https://github.com/szczurek-lab/SIEVE. DataFilter for selecting candidate variant sites is available at https://github.com/szczureklab/DataFilter. The simulator is hosted at https://github.com/szczurek-lab/SIEVE_simulator, and the reproducible benchmarking framework is available at https://github.com/szczurek-lab/SIEVE_benchmark_pipeline. The scripts for generating all figures in this paper are hosted at https://github.com/szczurek-lab/SIEVE_analysis. All aforementioned code are freely accessible under a GNU General Public License v3.0 license.

## Acknowledgments

We thank Dr. Timothy Vaughan for valuable instructions on package development for BEAST 2. This project has received funding from the European Union’s Horizon 2020 research and innovation programme under the Marie Sklodowska-Curie grant agreement No. 766030. E.S. acknowledges the support from the Polish National Science Centre SONATA BIS grant No. 2020/38/E/NZ2/00305. D.P. was supported by the European Research Council (ERC-617457-PHYLOCANCER), the Spanish Ministry of Science and Innovation (PID2019-106247GB-I00), and Xunta de Galicia.

## Author contributions

S.K. and E.S. conceived the SIEVE model - with input and feedback from J.K., N.BE. and D.P. S.K. implemented the model, performed all model performance analysis and generated all figures. S.PL., D.C. and D.P. performed the CRC28 scDNA-seq experiment. N.BO., M.V. and J.A. processed the scDNA-seq datasets. S.K. and E.S. wrote the manuscript with critical comments and input from all the co-authors. E.S. supervised the study.

## Extended data

**Extended Data Fig. 1:**
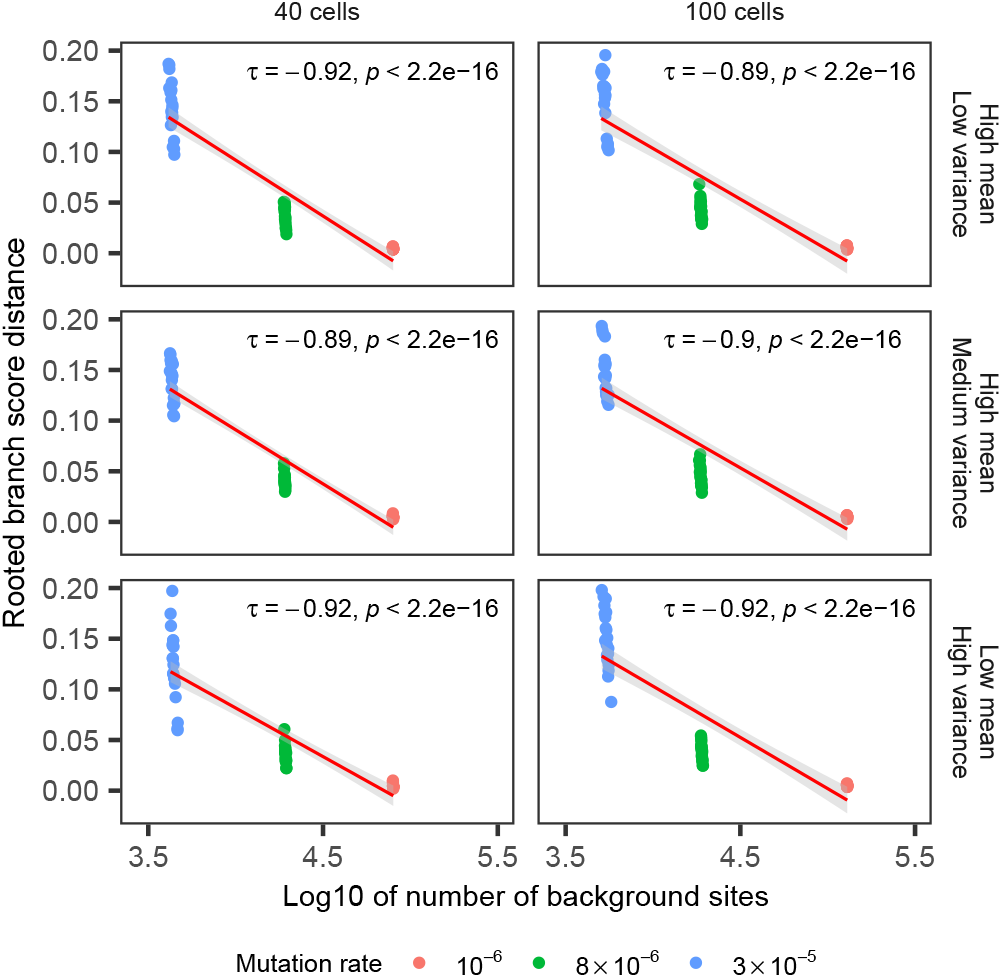
Correlation plot of the rooted BS distance against the number of background sites in log10 scale. Varying are the number of cells and the coverage quality. Rooted BS distance data points are coloured by the corresponding mutation rates. Kendall is the method for computing the correlation coefficient *τ*, which is invariant to the log transformation of the number of background sites. We choose 0.01 as the significance threshold.

**Extended Data Fig. 2:**
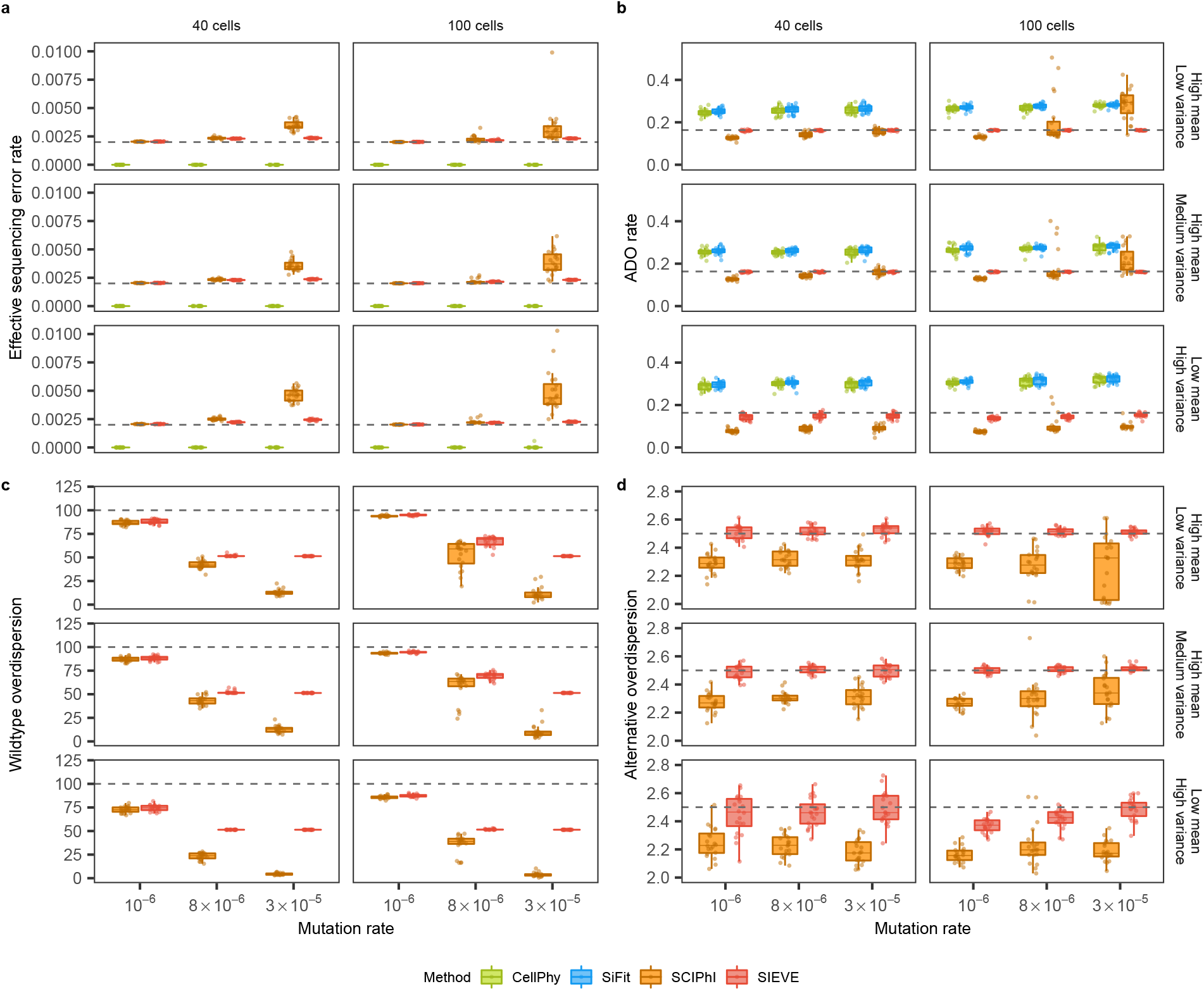
Additional benchmarking results of the SIEVE model regarding parameter estimates. Each simulation is repeated *n* = 20 times with each repetition denoted by coloured dots. The grey dashed lines represent the ground truth used to generate the simulated data. **a-d**, Box plots of parameter estimation accuracy for four important parameters in the model of raw read counts (Methods): effective sequencing error rate (**a**), ADO rate (**b**), wildtype overdispersion (**c**) and alternative overdispersion (**d**).

**Extended Data Fig. 3:**
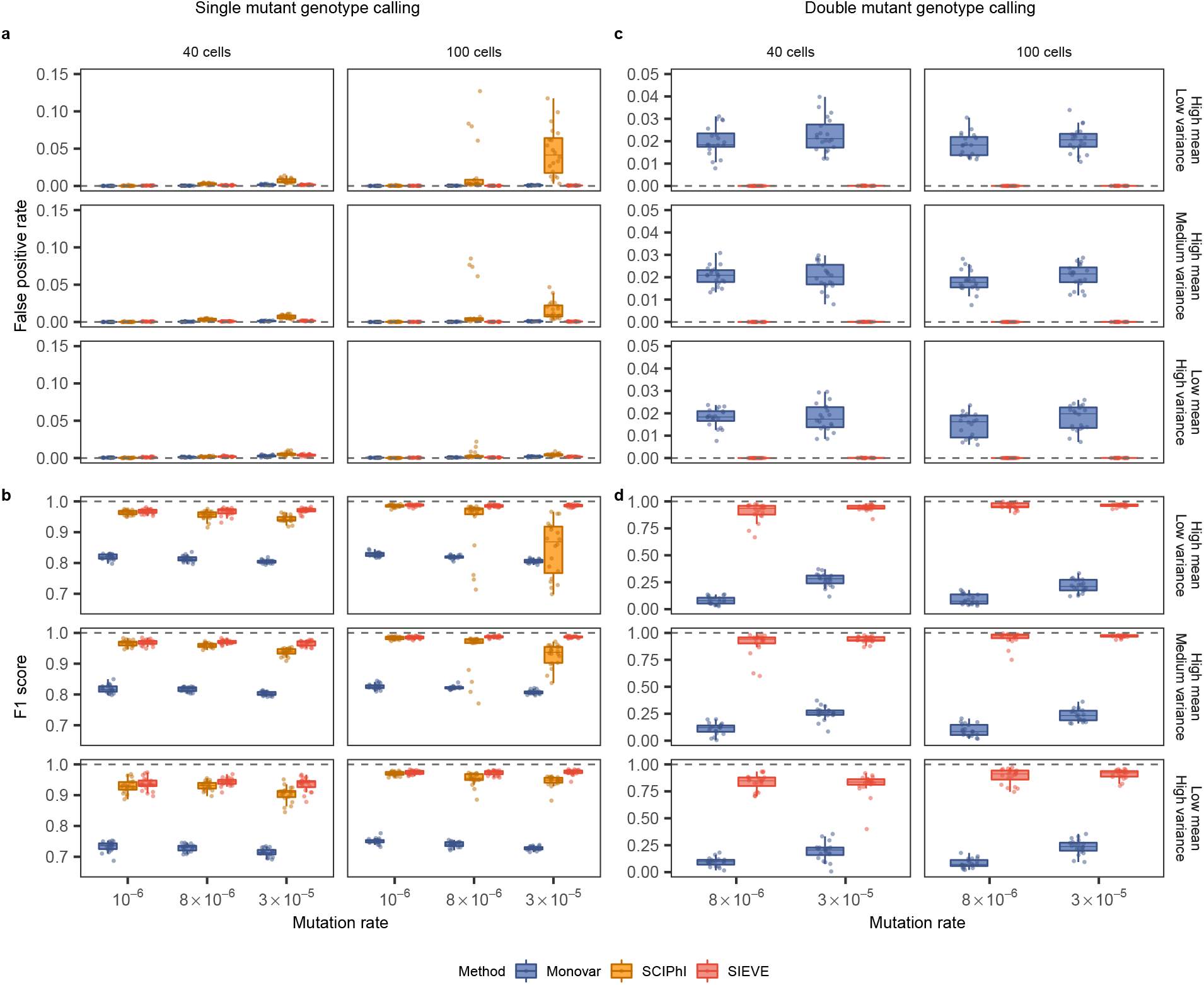
Additional benchmarking results of the SIEVE model regarding variant calling. Each simulation is repeated *n* = 20 times with each repetition denoted by coloured dots. The grey dashed lines represent the optimal values of each metric. **a-b**, Box plots of the single mutant genotype calling results measured further by the fraction of false positives in the ground truth negatives, i.e., the sum of false positives and true negatives, (false positive rate, **a**) and the harmonic mean of recall and precision (F1 score, **b**). **c-d**, Box plots of the double mutant genotype calling results measured further by false positive rate (**c**) and F1 score (**d**), where the variant calling results when mutation rate is 10^−6^ are omitted as very few double mutant genotypes are generated (less than 0.1%).

**Extended Data Fig. 4:**
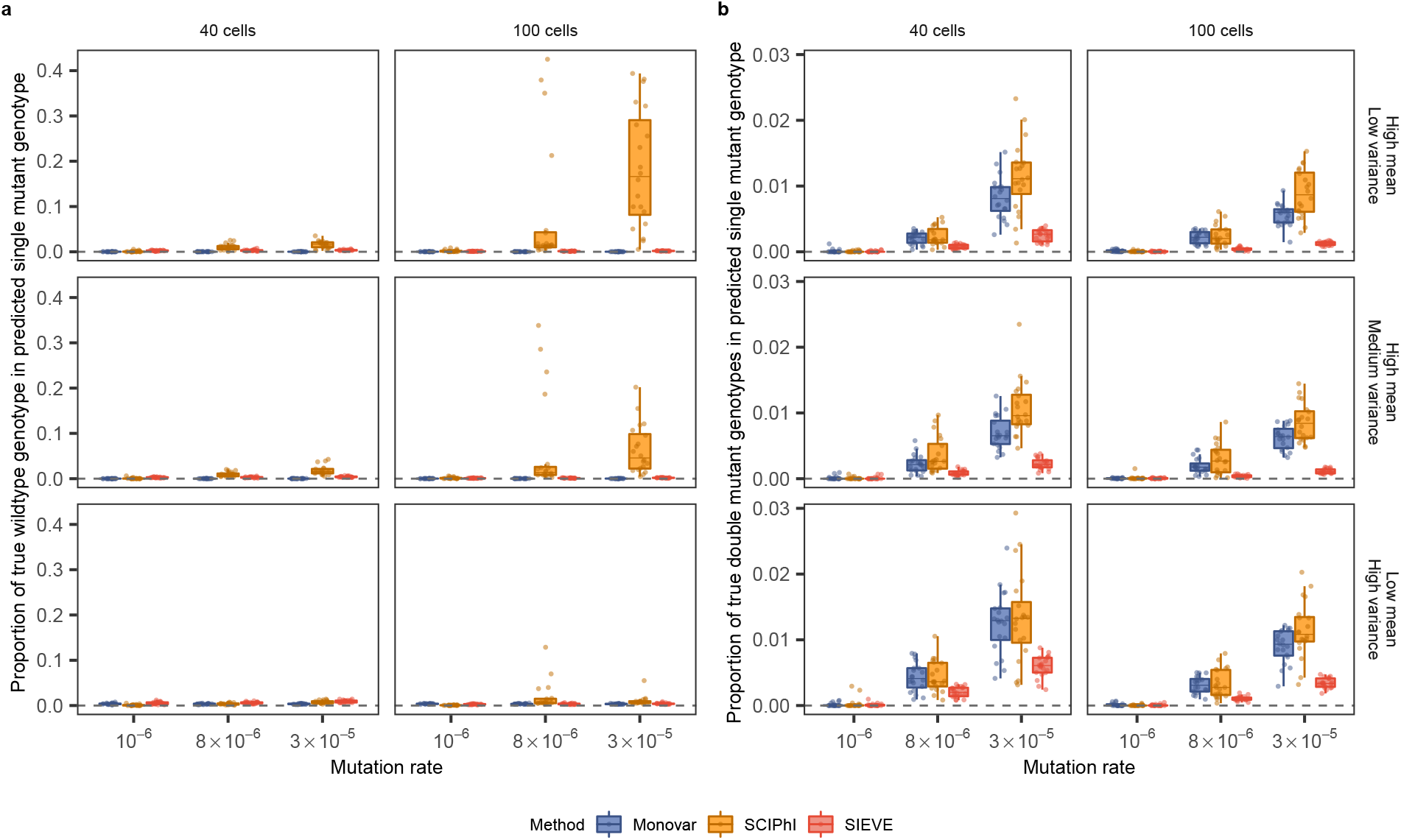
Types of false positives in single mutant genotype calling. The grey dashed lines represent the optimal proportions of each type. **a-b**, Box plots of the types of false positives in single mutant genotype calling, including the proportion of true wildtype (**a**) and true double mutant genotype (**b**). For single mutant genotype calling, the sum of the precision, the proportion of true wildtype and the proportion of true double mutant genotype is 1.

**Extended Data Fig. 5:**
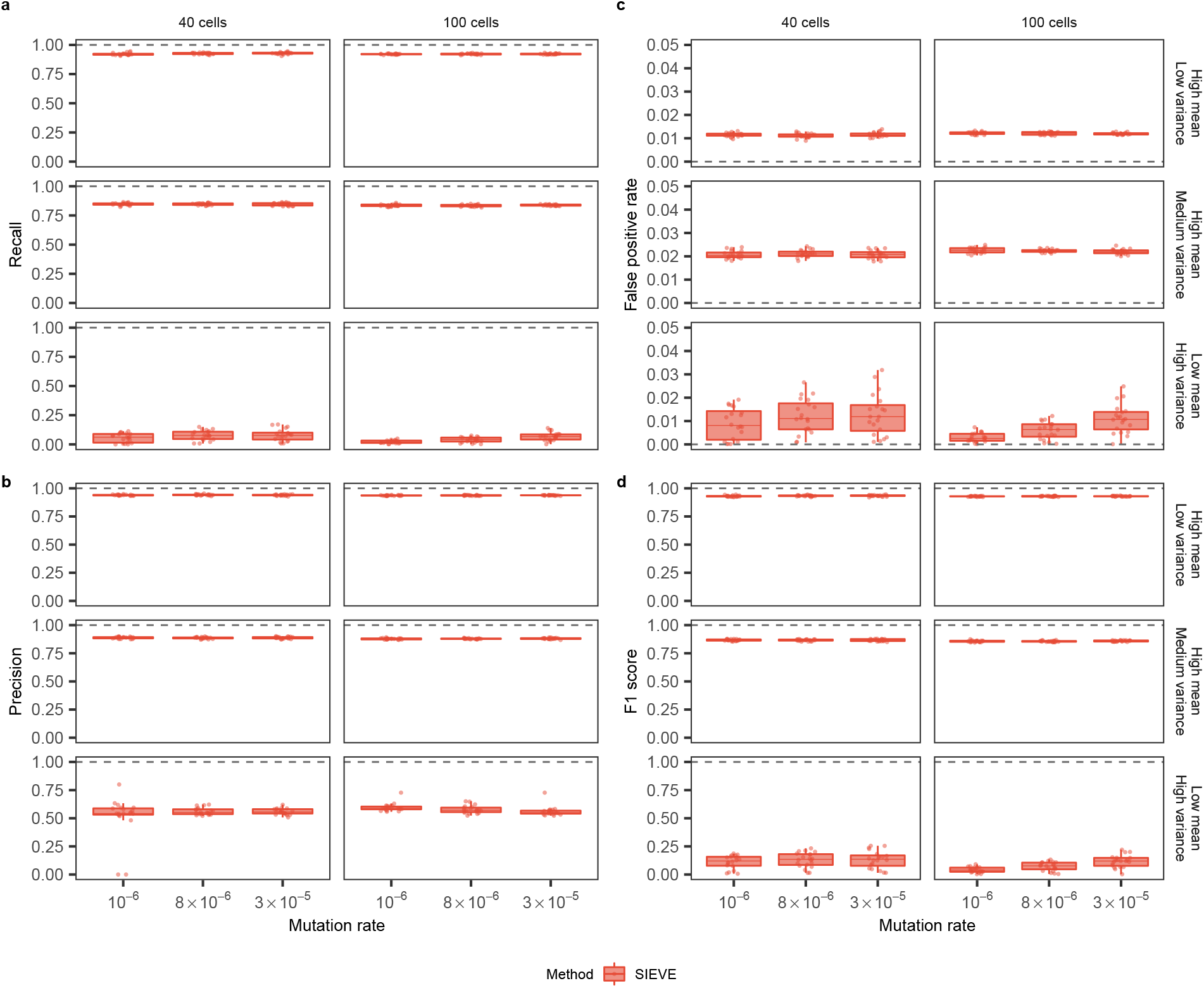
Benchmarking results of the SIEVE model regarding ADO calling. Each simulation is repeated *n* = 20 times with each repetition denoted by coloured dots. The grey dashed lines represent the optimal values of each metric. **a-d**, Box plots of the ADO calling results measured in recall (**a**), precision (**b**), false positive rate (**c**) and F1 score (**d**).

**Extended Data Fig. 6:**
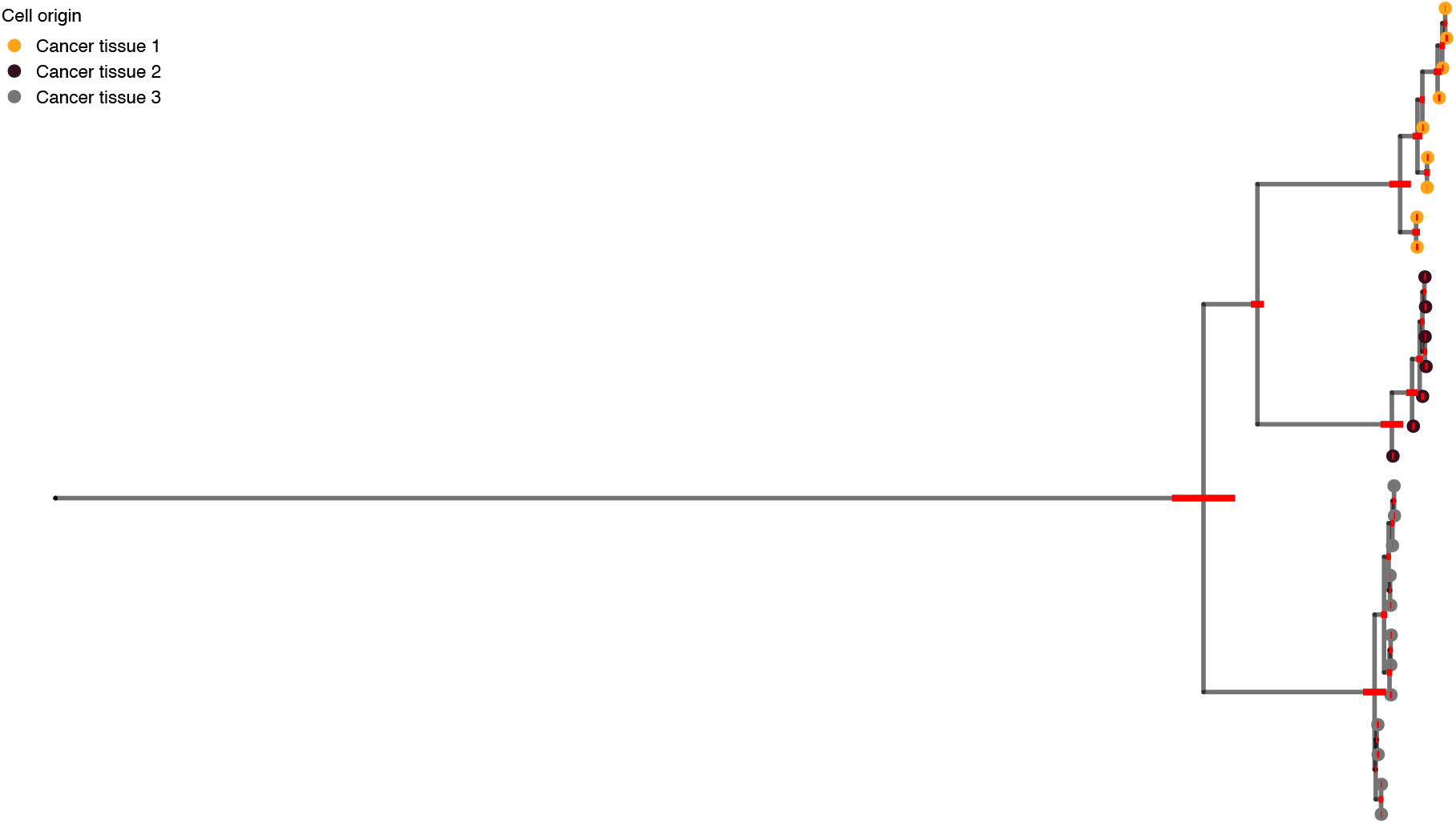
Illustration of branch lengths of the phylogenetic tree inferred from CRC28 by SIEVE. Shown is exactly the same tree as in Fig. 3, except that cell names, subclone posterior probabilities and gene annotations are removed and no branches are folded. Red bars annotated to internal nodes except the root are the 95% highest posterior density (HPD) intervals of the corresponding branch lengths.

**Extended Data Fig. 7:**
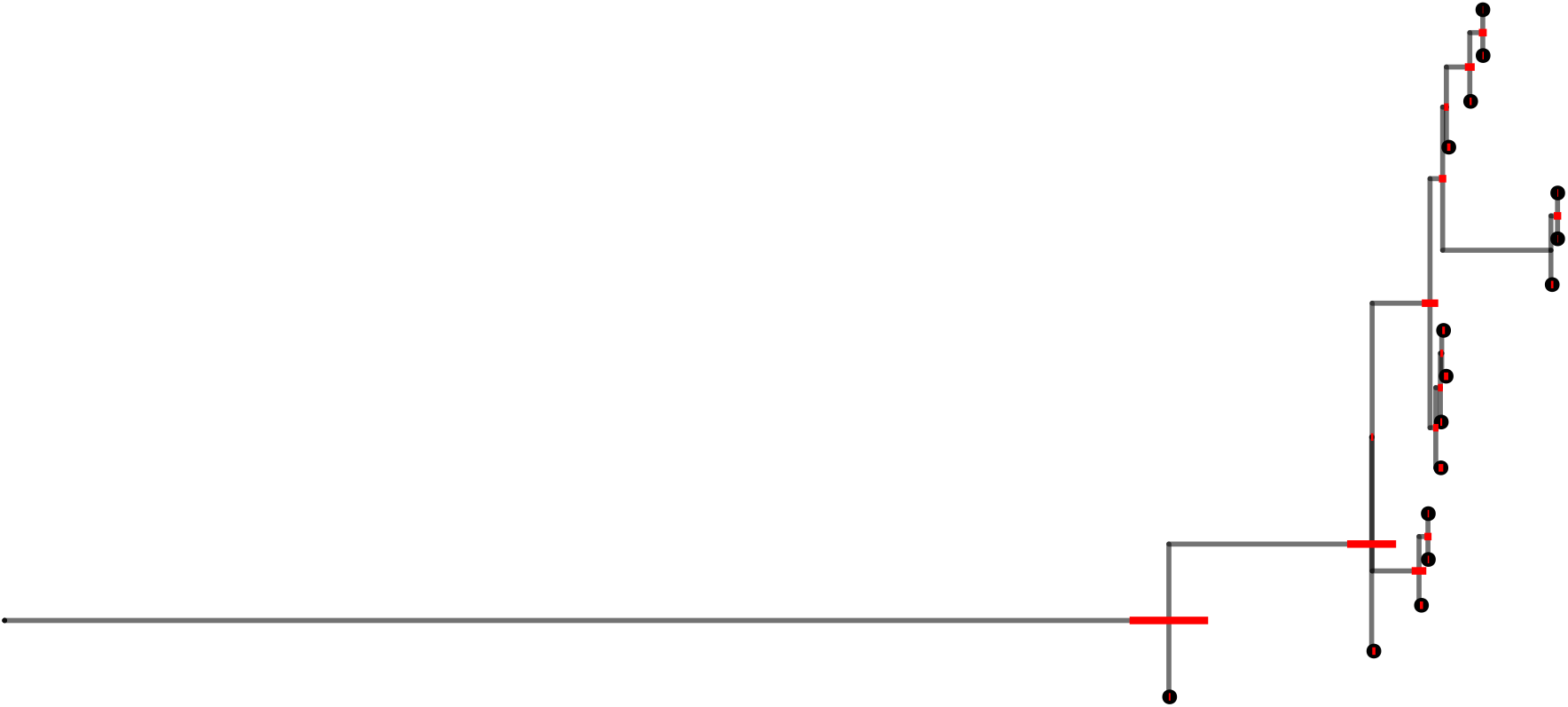
Illustration of branch lengths of the phylogenetic tree inferred from TNBC16 [40] by SIEVE. Shown is exactly the same tree as in Fig. 4, except that cell names, subclone posterior probabilities and gene annotations are removed and no branches are folded. Red bars annotated to internal nodes except the root are the 95% HPD intervals of the corresponding branch lengths.

**Extended Data Fig. 8:**
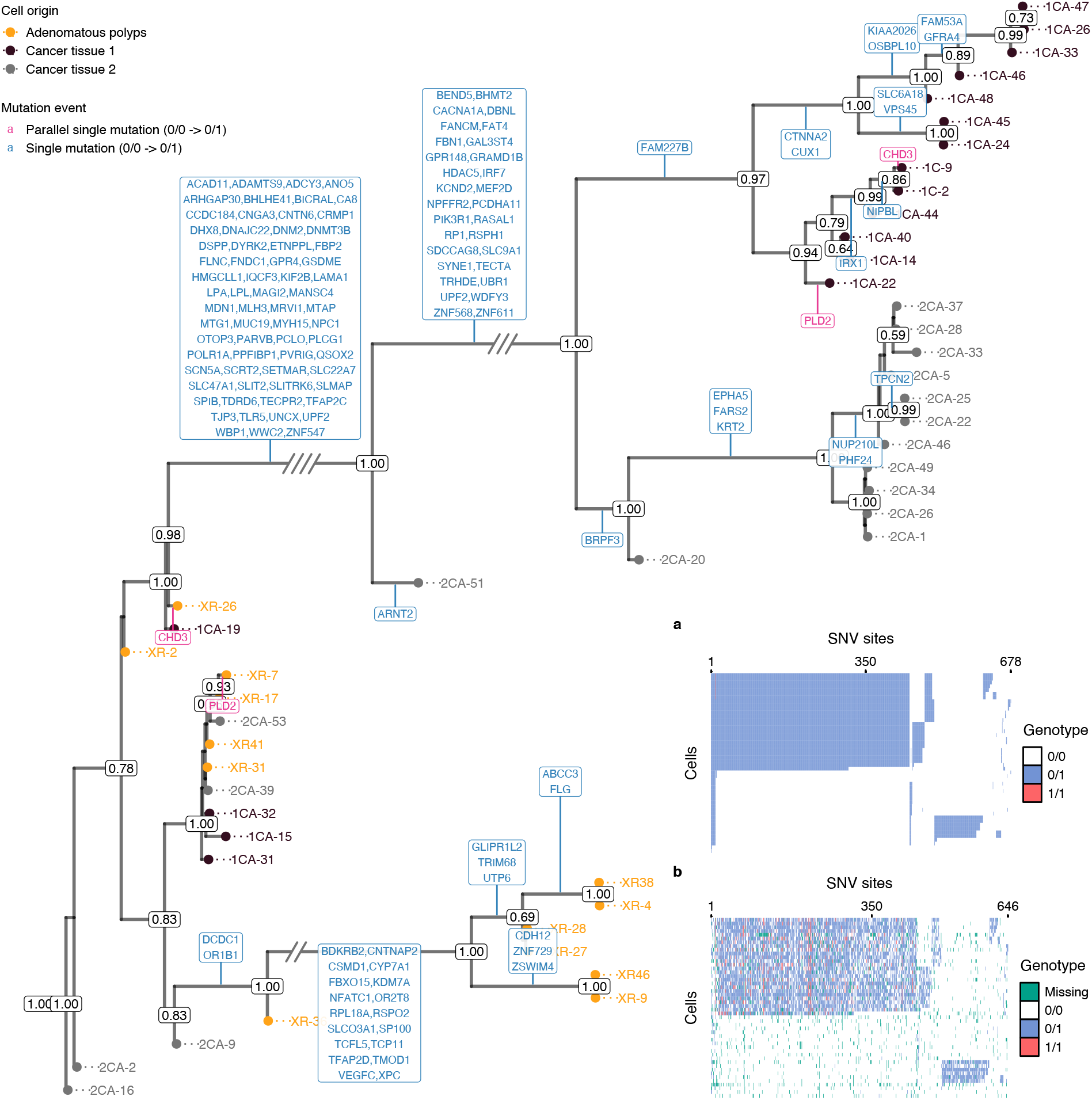
Results of phylogenetic inference and variant calling for CRC48 [41] dataset. Shown is SIEVE’s maximum clade credibility tree. Three exceptionally long branches are folded with the number of slashes proportional to the branch lengths. Cell names are annotated to the leaves of the tree, coloured by the corresponding biopsies. The numbers at each node represent the posterior probabilities (threshold *p >* 0.5). At each branch, non-synonymous mutations are depicted in different colours including single mutations in blue and parallel single mutations in pink. **a-b**, Variant calling heatmap for SIEVE (**a**) and Monovar (**b**). Listed in the legend are the categories of predicted genotypes by each method. Cells in the row are in the same order as that of leaves in the phylogenetic tree.

**Extended Data Fig. 9:**
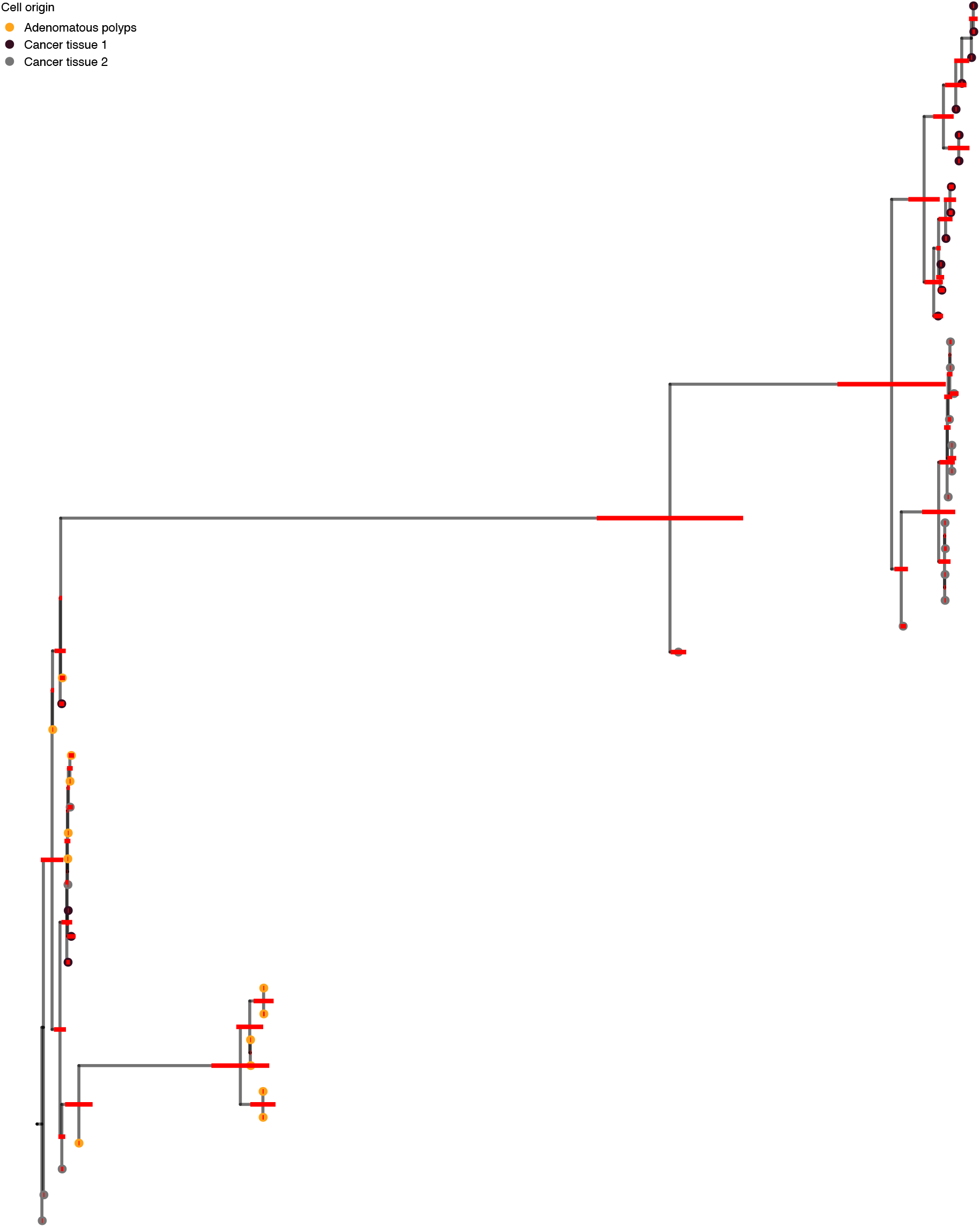
Illustration of branch lengths of the phylogenetic tree inferred from CRC48 [41] by SIEVE. Shown is exactly the same tree as in Extended Data Fig. 8, except that cell names, subclone posterior probabilities and gene annotations are removed and no branches are folded. Red bars annotated to internal nodes except the root are the 95% HPD intervals of the corresponding branch lengths.

**Extended Data Table 1:**
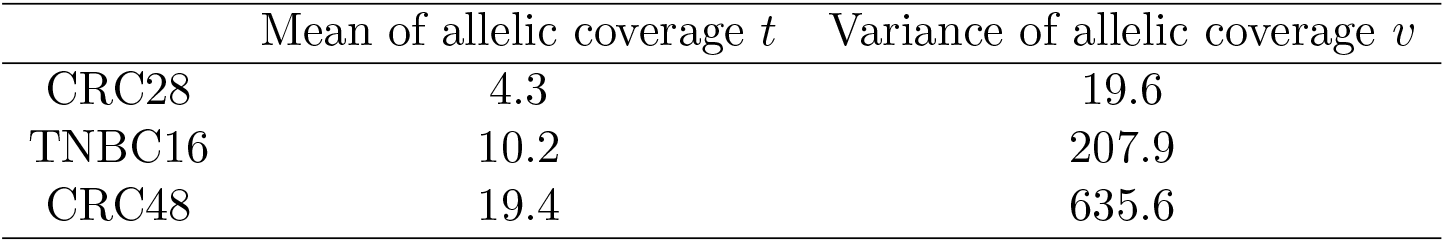
Inferred mean and variance of allelic coverage for real datasets.

| | Mean of allelic coverage $t$ | Variance of allelic coverage $v$ |
| --- | --- | --- |
| CRC28 | 4.3 | 19.6 |
| TNBC16 | 10.2 | 207.9 |
| CRC48 | 19.4 | 635.6 |

