## Supplementary Information for "SIEVE: joint inference of single-nucleotide variants and cell phylogeny from single-cell DNA sequencing data"

Senbai Kang[1], Nico Borgsmüller[2,3], Monica Valecha[4,5], Jack Kuipers[2,3], Joao Alves[4,5], Sonia Prado-López[4,5,6], Débora Chantada[7], Niko Beerenwinkel[2,3], David Posada[4,5,8], and Ewa Szczurek[1,*]

[1] *Faculty of Mathematics, Informatics and Mechanics, University of Warsaw, Warsaw, Poland*

[2] *Department of Biosystems Science and Engineering, ETH Zurich, 4058 Basel, Switzerland*

[3] *SIB Swiss Institute of Bioinformatics, 4058 Basel, Switzerland*

[4] *CINBIO, Universidade de Vigo, 36310 Vigo, Spain*

[5] *Galicia Sur Health Research Institute (IIS Galicia Sur), SERGAS-UVIGO*

[6] *Institute of Solid State Electronics E362, Technische Universität Wien, Austria*

[7] *Department of Pathology, Hospital Álvaro Cunqueiro, Vigo, Spain*

[8] *Department of Biochemistry, Genetics, and Immunology, Universidade de Vigo, 36310 Vigo, Spain*

### Supplementary table

| | | Missing | 0/0 | 0/1 | 1/1 | $1/1'$ |
|---|---|---|---|---|---|---|
| CRC28 | SIEVE | NA | 25.02% | 74.64% | 0.28% | 0.06% |
| | Monovar | 10.40% | 38.09% | 46.30% | 5.21% | NA |
| TNBC16 | SIEVE | NA | 15.54% | 75.11% | 9.30% | 0.05% |
| | Monovar | 10.63% | 32.92% | 41.58% | 14.87% | NA |
| CRC48 | SIEVE | NA | 59.48% | 40.50% | 0.02% | 0 |
| | Monovar | 4.53% | 69.41% | 24.13% | 1.93% | NA |

**Supplementary Table 1: Summary of fractions of predicted genotypes by SIEVE and Monovar for three analysed real datasets.** Entries marked with NA denote that the corresponding method does not call the specific genotype.

| | A/A | A/C | A/G | A/T | C/C | C/G | C/T | G/G | G/T | T/T |
|---|---|---|---|---|---|---|---|---|---|---|
| A/A | -1 | $1/3$ | $1/3$ | $1/3$ | 0 | 0 | 0 | 0 | 0 | 0 |
| A/C | $1/6$ | -1 | $1/6$ | $1/6$ | $1/6$ | $1/6$ | $1/6$ | 0 | 0 | 0 |
| A/G | $1/6$ | $1/6$ | -1 | $1/6$ | 0 | $1/6$ | 0 | $1/6$ | $1/6$ | 0 |
| A/T | $1/6$ | $1/6$ | $1/6$ | -1 | 0 | 0 | $1/6$ | 0 | $1/6$ | $1/6$ |
| C/C | 0 | $1/3$ | 0 | 0 | -1 | $1/3$ | $1/3$ | 0 | 0 | 0 |
| C/G | 0 | $1/6$ | $1/6$ | 0 | $1/6$ | -1 | $1/6$ | $1/6$ | $1/6$ | 0 |
| C/T | 0 | $1/6$ | 0 | $1/6$ | $1/6$ | $1/6$ | -1 | 0 | $1/6$ | $1/6$ |
| G/G | 0 | 0 | $1/3$ | 0 | 0 | $1/3$ | 0 | -1 | $1/3$ | 0 |
| G/T | 0 | 0 | $1/6$ | $1/6$ | 0 | $1/6$ | $1/6$ | $1/6$ | -1 | $1/6$ |
| T/T | 0 | 0 | 0 | $1/3$ | 0 | 0 | $1/3$ | 0 | $1/3$ | -1 |

**Supplementary Table 2: Evolutionary rate matrix used in the simulator to generate the simulated data. Genotypes are encoded with nucleotides rather than numbers.**

### Supplementary note

### Commands for data preprocessing

Similar preprocessing procedure was done for all real datasets. TNBC16 dataset is shown as an example below.

File containing raw sequencing data information:

```
### acc-cell.txt (A text file containing cell sra ids for download and \
corresponding cell names)

SRR1163012      a1
SRR1163013      a2
SRR1163019      a3
SRR1163026      a4
SRR1163027      a5
SRR1163034      a6
SRR1163035      a7
SRR1163043      a8
SRR1163053      h1
SRR1163070      h2
SRR1163074      h3
SRR1163083      h4
SRR1163084      h5
SRR1163091      h6
SRR1163095      h7
SRR1163148      h8
SRR1163149      TNBC_n1
SRR1163150      TNBC_n2
SRR1163151      TNBC_n3
SRR1163152      TNBC_n4
SRR1163153      TNBC_n5
SRR1163154      TNBC_n6
SRR1163155      TNBC_n7
```

```
SRR1163156      TNBC_n8
SRR1163157      TNBC_n9
SRR1163158      TNBC_n10
SRR1163159      TNBC_n11
SRR1163160      TNBC_n12
SRR1163161      TNBC_n13
SRR1163162      TNBC_n14
SRR1163163      TNBC_n15
SRR1163164      TNBC_n16
SRR1163508      TNBC_Pop_Normal
SRR1298936      TNBC_Pop_Tumor
```

Other files or constants needed for preprocessing:

```
REF=hs37d5.fa
DBSNP=dbsnp_138.b37.vcf
INDELS=Mills_and_1000G_gold_standard.indels.b37.vcf
INDELS2=1000G_phase1.indels.b37.vcf
### $SLURM_ARRAY_TASK_ID - chromosomes.
SLURM_ARRAY_TASK_ID = 1-22
```

**Downloading Data according to SRA using SRA-Toolkit.**

```
module load sra-toolkit/2.9.2-centos_linux64
cat acc-cell.txt | while read -r sra_id cellname;do
fastq-dump --split-files $sra_id
done
```

**Fastq Quality Control using CutAdapt.**

```
module load gcccore/6.4.0 cutadapt/1.18-python-3.7.0
cat acc-cell.txt | while read -r sra_id cellname;do
cutadapt --minimum-length 70 \
        -a AGATCGGAAGAGCACACGTCTGAACTCCAGTCACNNNNNNATCTCGTATGCCGTCTTCTGCTTG  \
        -o Processing/${sra_id}.trimmed_1.fastq.gz \
```

```
        Raw_Data/${sra_id}_1.fastq.gz > ${sra_id}_Cutadapt.txt
done
```

**Alignment using BWA.**

```
module load gcccore/6.4.0 cutadapt/1.18-python-3.7.0
module load gcc/6.4.0 bwa/0.7.17
module load picard/2.18.14

cat acc-cell.txt | while read -r sra_id cellname;do

ID=${cellname}
SM=$(echo ${cellname} | cut -d "_" -f1)
PL=$(echo "ILLUMINA")
LB=$(echo "KAPA")
PU=‘zcat Raw_Data/${cellname}_1.fastq.gz | head -n1 | \
        sed ’s/[:].*//’ | sed ’s/@//’ | sed ’s/ /_/’‘
echo "SAMPLE: "${cellname}" ID: "${ID}" SM: "${SM}
RG="@RG\\tID:${ID}\\tSM:${SM}\\tPL:${PL}\\tLB:${LB}\\tPU:${PU}"

bwa mem -t 10 \
        -R ${RG} \
        $REF \
        ./Processing/${sra_id}.trimmed_1.fastq.gz \
        > Processing/${cellname}.sam

java -Xmx18g -jar $EBROOTPICARD/picard.jar SortSam \
        I=Processing/${cellname}.sam \
        TMP_DIR=Processing/ \
        O=Processing/${cellname}.sorted.bam \
        CREATE_INDEX=true \
        SORT_ORDER=coordinate
done
```

**Mark Duplicates using Picard Tools.**

```
module load picard/2.18.14
cat acc-cell.txt | while read -r sra_id cellname;do
java -Xmx35g -jar $EBROOTPICARD/picard.jar MarkDuplicates \
        I=Processing/$cellname.sorted.bam \
        TMP_DIR=Processing/ \
        O=Processing/${cellname}.dedup.bam \
        CREATE_INDEX=true \
        VALIDATION_STRINGENCY=LENIENT \
        M=Processing/Duplicates_${cellname}.txt
done
```

**Realignment using GATK.**

```
module load gatk/3.7-0-gcfedb67

sample_bams=$(ls Processing/*.dedup.bam)
bams_in=$(echo $sample_bams | sed ’s/ / -I /g’)
echo $sample_bams |sed ’s/ /\n/g’ | sed ’s/.dedup.bam//g’ | \
        awk -v chr=$SLURM_ARRAY_TASK_ID \
        ’{print $0".dedup.bam\t"$0".real."chr".bam"}’ | \
        sed ’s/Processing\///’ > W32.$SLURM_ARRAY_TASK_ID.map

java -Djava.io.tmpdir=Processing/ -Xmx25G \
        -jar $EBROOTGATK/GenomeAnalysisTK.jar \
        -T RealignerTargetCreator \
        -I $bams_in \
        -o W32.$SLURM_ARRAY_TASK_ID.intervals \
        -R $REF \
        -known $INDELS \
        -known $INDELS2 \
        -L $SLURM_ARRAY_TASK_ID

```

```
java -Djava.io.tmpdir=Processing/ -Xmx25G \
        -jar $EBROOTGATK/GenomeAnalysisTK.jar \
        -T IndelRealigner \
        -known $INDELS \
        -known $INDELS2 \
        -I $bams_in \
        -R $REF \
        -targetIntervals W32.$SLURM_ARRAY_TASK_ID.intervals \
        -L $SLURM_ARRAY_TASK_ID \
        --nWayOut W32.$SLURM_ARRAY_TASK_ID.map \
        --maxReadsForRealignment 1000000
```

**Recalibration using GATK.**

```
module load gatk/4.0.10.0
cat acc-cell.txt | while read -r sra_id cellname;do
gatk --java-options "-Xmx24G -Djava.io.tmpdir=Processing/" BaseRecalibrator \
        -I Processing/$cellname.real.$SLURM_ARRAY_TASK_ID.bam \
        -O Processing/$cellname.recal.$SLURM_ARRAY_TASK_ID.table \
        -R $REF \
        --known-sites $DBSNP \
        --known-sites $INDELS

gatk --java-options "-Xmx24G -Djava.io.tmpdir=Processing/" ApplyBQSR \
        -R $REF \
        -I Processing/$cellname.real.$SLURM_ARRAY_TASK_ID.bam \
        --bqsr Processing/$cellname.recal.$SLURM_ARRAY_TASK_ID.table \
        -O Processing/$cellname.recal.$SLURM_ARRAY_TASK_ID.bam
done
```

**Bam to mpileup using samtools.**

```
module load samtools/1.9
ls ../Processing/*.recal.$SLURM_ARRAY_TASK_ID.bam | \
```

```
        grep -v "Tumor" > bampath.$SLURM_ARRAY_TASK_ID.txt
samtools mpileup --no-BAQ --min-BQ 13 --max-depth 10000 --min-MQ 40 \
        -r $SLURM_ARRAY_TASK_ID -b bampath.$SLURM_ARRAY_TASK_ID.txt \
        -f $REF -o W32.$SLURM_ARRAY_TASK_ID.mpileup
```
